# The hippocampus enables abstract structure learning without reward

**DOI:** 10.64898/2026.02.14.705916

**Authors:** Adedamola Onih, Xinran Shen, Lida Pentousi, Vezha Boboeva, Athena Akrami

## Abstract

Statistical learning (SL) allows organisms to infer latent structure from sensory input without instruction, feedback, or reward, yet how the brain accomplishes such abstract, unsupervised learning remains unknown. Here we show that mice, like humans, rapidly acquire multiple forms of statistical structure, including event frequency, sequence identity, and abstract structural rules, and that the hippocampus is essential for this capacity. Pupil dynamics provided a cross-species, implicit readout of expectation formation, revealing spontaneous sensitivity to these regularities during passive listening. In mice, pharmacological and temporally precise optogenetic inactivation of dorsal CA1 abolished all learning-related pupil signatures without affecting baseline pupil size, target-evoked responses, or task performance, demonstrating a causal requirement for the hippocampus in forming and updating internal models of sensory structure. High-density recordings further revealed that dCA1 ensembles track evolving statistical contexts while dynamically reorganising population activity into subspaces that separately encode sensory features and abstract rules, enabling generalisation across distinct but structurally equivalent sequences. Together, these results identify the hippocampus as a critical neural substrate for latent abstract structure learning and offer a mechanistic account of how internal models emerge from unsupervised experience.

## Introduction

In a world filled with sensory complexity and uncertainty, organisms must learn not only from rewards and outcomes but also by detecting structure embedded in ongoing experience. This ability to acquire knowledge without instruction, feedback, or reinforcement, termed statistical learning (SL), is essential for anticipating events, segmenting experience, and generalising across contexts (Fiser et al., 2010). Because explicit rewards and instructions are sparse in natural environments, much of real-world learning is unsupervised i.e. incidental and reward-independent (Bröker et al., 2024). SL is conserved across species and sensory modalities and supports core cognitive functions including perception, decision-making, and language acquisition (Schapiro et al., 2014; Schapiro & Turk-Browne, 2015). By extracting latent structure from experience without external guidance or reward, SL enables the brain to form internal models that guide flexible, predictive behaviour.

Early work showed that human infants could detect regularities in streams of artificial syllables, allowing them to segment word-like units from continuous input (Saffran et al., 1996). Subsequent work demonstrated that SL extends across sensory (Barascud et al., 2016; Conway & Christiansen, 2005; Creel et al., 2004; Fiser & Aslin, 2002; Henin et al., 2021; Lengyel et al., 2019; Turk-Browne et al., 2005, 2009) and throughout the lifespan, supporting rapid acquisition of structure ranging from simple co-occurrences to abstract rules that generalise across surface variations (Bulf et al., 2011; Saffran et al., 1999; Thiessen, 2010; Wetzel et al., 2016).

Together, these findings establish SL as a general property of cognition and raise a central question: how does the brain rapidly construct internal models of abstract structure from unlabelled sensory experience?

Evidence from human lesion studies implicates the medial temporal lobe, and the hippocampus in particular, in SL (Aljishi et al., 2023; Covington et al., 2018; Schapiro et al., 2014). This has motivated theoretical accounts in which the hippocampus acts as a predictive engine, compressing regularities across experience into flexible representations for inference (Behrens et al., 2018; Buzsáki & Tingley, 2018; Fang et al., 2023; Greco et al., 2024; Koster et al., 2018; Levenstein et al., 2024; Penny et al., 2013; Raju et al., 2024; Stachenfeld et al., 2017; Tarder-Stoll et al., 2024). Neuroimaging and electrophysiology in humans support this view (Aljishi et al., 2023; Hannula & Greene, 2012; Henin et al., 2021; Hindy et al., 2016; Kok et al., 2019; Kumaran & Maguire, 2009; Schapiro et al., 2012), but these approaches lack the temporal resolution and causal leverage needed to resolve the circuit-level computations underlying SL. A mechanistic understanding of how hippocampal circuits build predictive models from unsupervised experience therefore remains missing.

Animal models provide this mechanistic access, yet suitable paradigms for SL have been limited. Most rodent learning tasks probe reinforcement learning or fear conditioning, where behaviour is shaped by explicit cues, rewards, or simple associations (Menichini et al., 2025; Xiao et al., 2018; Yi et al., 2022). In parallel, a rich, cross-species literature on predictive coding shows that rodents and primates detect local and global structures in passive settings, including mismatch negativity, stimulus-specific adaptation, and higher-order deviant detection (Bekinschtein et al., 2009; Libby & Buschman, 2021; Luo et al., 2025; Meyer & Olson, 2011; Parras et al., 2017; Ulanovsky et al., 2003). While these paradigms reveal sensory and motor prediction-error signals, they primarily probe short-term expectation violations and do not address how animals infer latent structure that accumulates gradually across repeated, temporally separated exposures—a central property of statistical learning. Moreover, since these passive paradigms typically lack a behavioural or physiological measure of acquired knowledge, it remains unclear whether structure is actually learned. Sequence learning has been demonstrated in birds and primates (Abe & Watanabe, 2011; Fehér et al., 2017; Gentner et al., 2006; Milne et al., 2021), but analogous reinforcement-free rodent paradigms with an interpretable readout of learning have been lacking, preventing use of modern circuit-level tools to investigate the neural basis of SL.

To address this gap, we developed a novel reinforcement-free behavioural paradigm for studying SL in mice and humans under matched conditions. Subjects performed a simple auditory cover task in form of detection of a broadband target sound, while task-irrelevant tone sequences, governed by latent statistical structure, were embedded in the background stream. To track learning without explicit report, we used pupil dilation as a non-verbal readout of surprise and expectation (Alamia et al., 2019; Friedman et al., 1973; Liao et al., 2016; Qiyuan et al., 1985; Quirins et al., 2018; Zhao et al., 2019a). Although pupil size is modulated by arousal and attention (Aston-Jones & Cohen, 2005; Stanners et al., 1979), converging evidence shows that phasic pupil responses scale with expectation violations (Joshi & Gold, 2020; Zhao et al., 2019b). Critically, surprise can only occur if expectations have been formed through learning; in the absence of learning, violations are not surprising. We therefore leveraged pupil dynamics as an implicit measure of expectation formation, while explicitly controlling for baseline arousal, sensory responsiveness, or task engagement.

Importantly, our paradigm engages computations foundational to statistical learning: binding and segmentation of multi-tone sequences embedded in a continuous background stream to extract sequence identity; integration across tens of seconds and many trials to infer event statistics by linking temporally separated exposures; and inference of latent abstract structure shared across physically distinct sequences, enabling generalisation beyond surface features. These operations, i.e. long-range temporal integration, flexible binding across discontinuous inputs, and abstraction over sensory detail, are strongly associated with hippocampal–entorhinal circuitry rather than primary sensory cortices (Aronov et al., 2017; Schapiro et al., 2012, 2014, 2016). By engaging precisely these computations, our task provides a mechanistic handle on SL that classical deviance-detection paradigms cannot access.

Using this framework, we show that mice, like humans, rapidly acquire multiple forms of statistical structure, including event frequency, pattern identity, and abstract structural rules, purely from incidental experience. We further demonstrate a causal role for dorsal CA1 (dCA1) in this process: pharmacological and temporally specific optogenetic inactivation abolished learning related pupil signatures while sparing auditory processing, baseline pupil dynamics, target-evoked responses, and task performance. High-density recordings revealed that dCA1 population activity simultaneously encoded sensory features and latent structure, dynamically factorising sensory information and abstract structure into orthogonal subspaces. This geometric organisation allows the hippocampus to represent the latent ‘rule’ independently of the specific sensory content, enabling the rapid generalisation we observed behaviourally.

Together, these results establish a rodent model of SL and provide the first causal, circuit-level evidence that the hippocampus, via dCA1, constructs abstract, generalisable internal models from unsupervised experience.

## Results

### Mice and humans rapidly extract sensory regularities without instruction or reinforcement

To investigate the neural basis of unsupervised statistical learning, we developed a novel cross-species paradigm in which structured, task-irrelevant auditory sequences were embedded sparsely and unpredictably within a continuous sound stream, while subjects performed a cover task. Mice were trained to lick in response to a broadband noise target (X) while ignoring structured tone sequences (e.g. ABCD) presented at random times (Fig. 1A, Fig.1B). Animals quickly learned to respond to the X stimulus and showed robust licking behaviour, responding selectively to the X stimulus (Fig. 1D, Fig. S1A). Performance, reaction time and lick rate in response to X were unaffected by the presence of pattern sequence (Fig. 1C, Fig.1D, Fig. S1C-D) and whether the sequence preceded, followed, or overlapped with the target, and regardless of its timing in the trial (Fig. S1E–G). Thus, animals remained engaged with the cover task, while the sequences were behaviourally irrelevant and temporally unpredictable.

**Figure 1.**
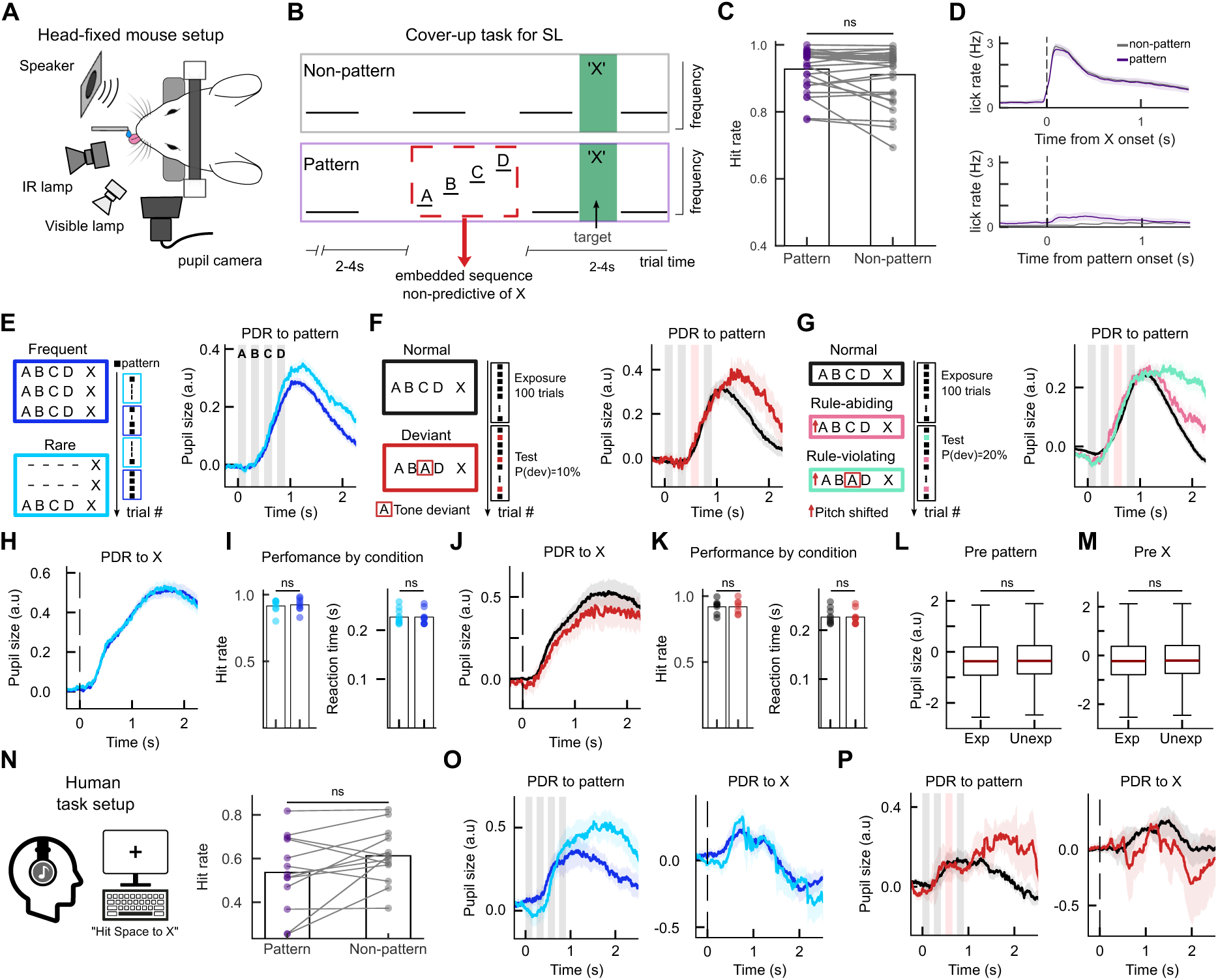
Pupil dynamics reveal spontaneous acquisition of statistical structure in mice and humans. **(A)** Schematic of the head-fixed behavioural setup **(B)** Auditory cover-up task structure. Mice lick in response to a broadband noise target (X, green) while ignoring embedded (task irrelevant) tone sequences. Dashed lines indicate repeating baseline tones; box highlights an example embedded sequence (ABCD). Timing of embedded sequence was independent of X onset. **(C)** Mean hit rate for target X across animals on trials with (pattern) versus without (non-pattern) embedded sequences. **(D)** Average lick rate aligned to X onset for pattern and non-pattern trials. Shaded areas indicate SEM across mice. **(E) Sensitivity to event rate.** Left: schematic of rare (10%) versus frequent (90%) sequence presentation block. Right: pupil dilation responses (PDR) aligned to pattern onset. Lines show mean across sessions; shaded error bands show SEM (n = 223 sessions from 8 mice). **(F) Sensitivity to sequence identity.** Similar to F, for Normal versus Deviant sequences (n = 28 sessions from 7 mice). **(G) Sensitivity to abstract structural rule.** Similar to E, for rule-conforming versus rule-violating deviants (n = 63 sessions from 6 mice). **(H)** Control PDR measures to x for rare vs frequent trials. Event rate does not modulate pupil in response to the target sound **(I)** Hit rate (left) and reaction time (right) for rare vs frequent trials. Event rate does not impact task performance. **(J-K)** similar to (H-I) for normal vs deviant trials. **(L)** Comparing the pre-pattern baseline pupil size for expected sequence events (frequent, normal) vs unexpected events (rare, deviant) **(M)** similar to (L) but for baseline pupil size before X appears. (H-M) together show that statistical manipulations in the task do not impact overall task engagement, global attention or arousal state. **(N-P) Human cross-species validation**. **(N)** Human behavioural performance (hit rates) on pattern versus non-pattern trials. **(O)** Human PDR for rare versus frequent sequences (n = 7 subjects) **(P)** Human PDR for Normal versus Deviant sequences (n = 5 subjects).

Critically, the statistical structure we probe is not available from any brief local stimulus history. Embedded sequences occurred at variable latencies within each trial (2–4 s after trial onset), and X occurred later and independently (5–10 s). This design forces learning to rely on integrating evidence across many discontinuous exposures over tens of seconds, rather than detecting immediate sensory irregularities.

To assess whether mice nonetheless learned the underlying structure of these sequences, we manipulated statistical regularities across three dimensions: event frequency, sequence identity, and abstract structural rules. In all cases, we used pupil dilation response (PDR) as a non-verbal, non-instrumental measure of surprise to unexpected auditory events.

### Sensitivity to event frequency

In a first manipulation, a given sequence (e.g. ABCD or see Fig. S2D for ABAD) was presented either frequently (90% of trials) or rarely (10% of trials) across alternating blocks. Despite being identical in acoustic content, rare sequences evoked significantly larger pupil dilations than the same sequences when frequent (shuffle-null permutation test, rare > frequent: p < 1×10⁻⁴; 10,000 iterations) (Fig. 1E), indicating that mice had acquired expectations about event rate.

Broadband noise targets (X), which were behaviourally relevant, and the onset of structured tone sequences, which were behaviourally irrelevant, both elicited large PDRs across sessions, consistent with their unpredictability in time. Within each trial, the embedded sequence and X could occur at variable latencies (2–4 s, and 5-10 s after trial onset, respectively), so both rare and frequent sequences were similarly ‘surprising’ at the within-trial timescales. Moreover, even in frequent blocks, the sequences remained sparse across trials (∼17 s mean inter-sequence interval, Fig. S2A), making their absolute timing hard to predict from any short window. Therefore, the reduced PDR for frequent versus rare sequences reflects learning at a slower timescale: mice extracted higher-order statistics about event rate across many discontinuous exposures, treating rare sequences as more surprising because they violated an across-trial estimate of probability, not because of when they occur within a trial.

Importantly, the reaction time and the lick response, as well as the PDR evoked by X did not differ across rare vs. frequent blocks (Fig. 1H-I). Similarly, the baseline pupil response remained the same across both blocks (Fig. 1M-L). Thus, statistical manipulations selectively modulated PDRs to the embedded sequences, without changing global arousal, attention, sensory responsiveness, or task engagement.

### Sensitivity to sequence identity

Next, we asked whether mice could detect violations of a learned sequence. Animals were first exposed to a consistent pattern (ABCD) for 100 trials (exposure phase) before presentation of a deviant pattern (ABAD) on 10% of subsequent trials (test phase). To isolate sequence violation from novelty, all sequences began with the same A and B tones, and no new sounds were introduced. Across all animals and sessions, deviant sequences evoked significantly larger pupil dilations than the normal sequence (shuffle-null permutation test, deviant > normal: p = < 1×10⁻⁴; 10,000 iterations) with the divergence emerging only after the third, where the deviant tone occurred (Fig. 1F). Notably, because sequences are separated by long, unpredictable gaps, detecting this identity violation requires binding tones into a coherent multi-tone chunk and comparing it to an internal model formed over many prior exposures, not merely responding to a transient change in sensory input. As in the event-rate manipulation, PDRs to the task-relevant target (X), the detection performance (i.e. the hit rate and the reaction time) and baseline pupil size remained stable across conditions (Fig. 1J-K), confirming that arousal changes were specific to sequence violations and independent of task engagement. As in the event-rate manipulation, this dissociation confirms that learning-related pupil effects were selective to sequence structure and independent of global arousal or sensory responsiveness.

To ensure that these effects were not idiosyncratic to specific tone combinations, we ran an additional experiment in which the same cohort of mice were exposed to ABAD serving as the normal sequence (for 100 trials) and ABCD as the deviant (on 10% of subsequent trials). The same pattern of results was observed, confirming that mice can learn arbitrary tone sequences and signal violations thereof (Fig. S2G-H).

### Sensitivity to abstract structure

Finally, we asked whether mice merely memorised specific tone transitions or inferred the latent abstract rule governing the sequences (e.g., ‘rising pitch’). To test for rule abstraction, we adapt the previous normal-deviant paradigm: after exposure to the standard rising sequence (ABCD_0_), animals were presented with two novel deviant types during the test phase: (1) a rule-conforming deviant (ABCD_1_) that used different frequencies but preserved the rising structure, and (2) a rule-violating (ABBA_1_), consisting of similar new tones, but violated the rising rule. Crucially, if mice relied on sensory memory, both new sequences should be equally surprising (equally novel). However, if they had learned the abstract “rising” rule, they should perceive ABCD_1_ as consistent with their internal model and ABBA_1_ as a violation. Both deviants evoked larger PDRs than to the original pattern, but consistent with rule learning, rule-violating deviants elicited significantly stronger pupil dilations than rule-conforming ones (shuffle-null permutation test, A_1_B_1_B_1_A_1_> A_1_B_1_C_1_D_1_: p = < 1×10⁻⁴; A_1_B_1_C_1_D_1_> normal: p = < 1×10⁻⁴; A_1_B_1_B_1_A_1_> normal: p = < 1×10⁻⁴; 10,000 iterations) (Fig. 1G). Thus, mice spontaneously generalised the learned structure to novel sensory inputs (“zero shot” transfer), implying a rule-level internal model rather than a stored template of specific tones.

The same paradigms in humans yielded similar effects: behavioural performance remained unaffected by the embedded structured sequence (Fig. 1N), while rare and deviant patterns produced significantly larger dilations than expected events (shuffle-null permutation test, rare > frequent: p < 1×10⁻⁴; deviant > normal: p < 0.006 10,000 iterations) (Fig. 1O, Fig.1P).

Together, pupil dilation responses provide a non-declarative window into expectation formation across extended timescales: both mice and humans rapidly and implicitly learned event frequency (requiring integration across many temporally separated exposures), sequence identity (requiring binding/segmentation across tones despite long inter-event gaps), and abstract rules (requiring generalisation beyond sensory features). Crucially, these effects were selective to the structured embedded sequences, meaning that baseline pupil size, pupil responses to the task-relevant target (X), hit rates and reaction times were unchanged across statistical manipulations (Fig. 1H-M). This demonstrates that changes in pupil dynamics reflect learning-dependent expectation violations rather than nonspecific modulation of arousal, attention, or task engagement.

### dCA1 is required for learning and updating statistical structure

Pupil responses showed that mice form expectations about auditory sequences and signal violations when the underlying statistics change (Fig.1). The statistical features manipulated here, i.e. (i) extracting event rate across tens of seconds with ∼17 s inter-sequence gaps, (ii) binding sparse multi-tone sequences to learn identity, and (iii) generalising abstract structure across sensory exemplars, require long-range temporal integration and mapping beyond sensory specifics; functions strongly associated with hippocampal circuitry and not readily accounted for by cortical sensory circuits. We therefore first asked, using a broad pharmacological manipulation, whether dorsal CA1 (dCA1) is necessary for this form of statistical learning, and then used temporally precise optogenetic silencing to pinpoint the time window during which dCA1 activity is required (Fig. 2).

**Figure 2.**
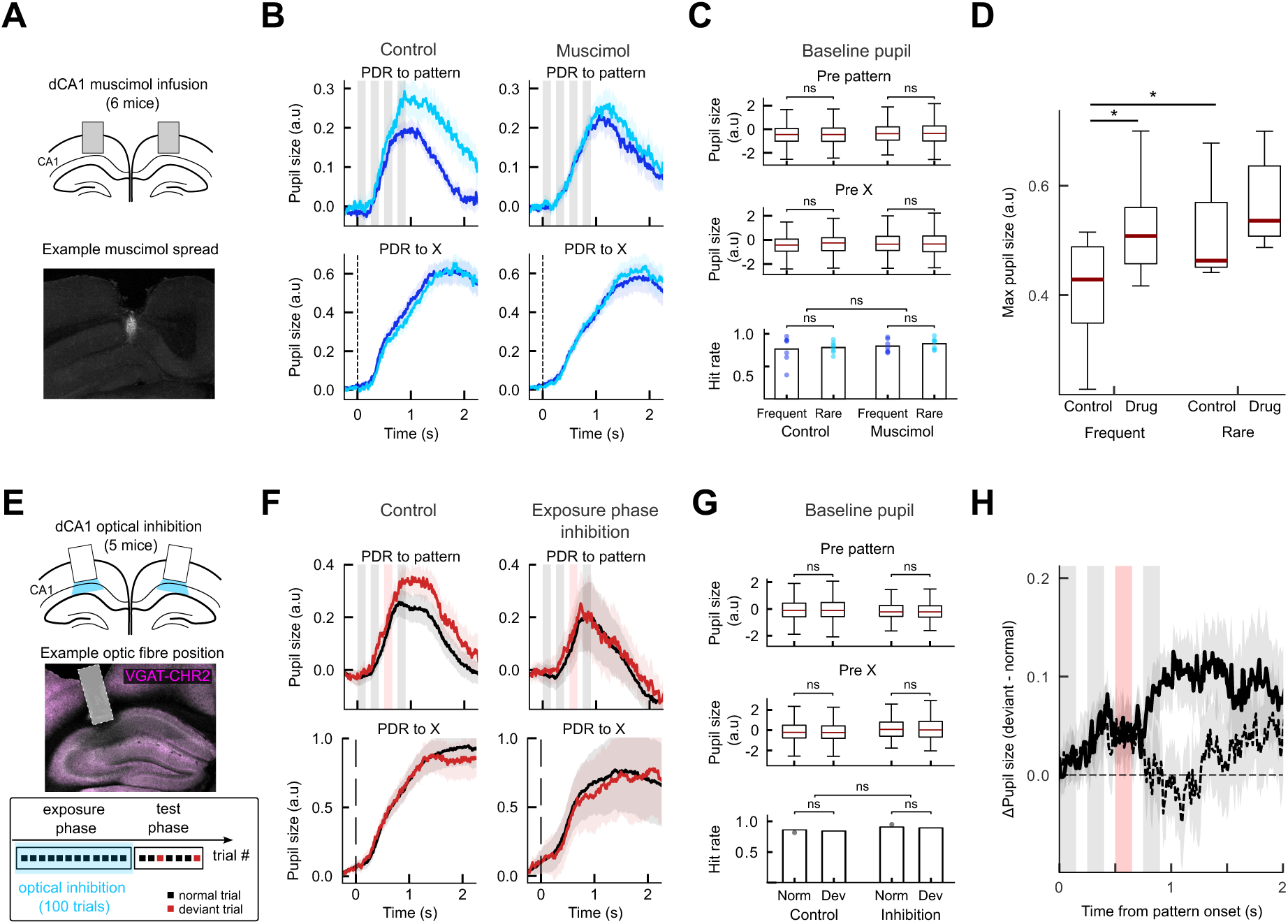
Dorsal CA1 is causally required for statistical learning independent of global arousal. (A) Schematic of bilateral cannula placement for muscimol infusion into dCA1 (top). Example histology image showing fluorescent muscimol spread in dorsal CA1 (bottom). Scale bar, 100 µm. (B) dCA1 silencing abolishes sensitivity to event frequency. Top: Pupil dilation responses (PDR) to rare (dark blue) vs. frequent (light blue) sequences in control (left) and muscimol (right) sessions. Bottom: PDR to the task-relevant target (X) is unaffected, confirming intact sensory responsiveness. Lines show mean across sessions; shaded bands show SEM. Gray shading marks tone presentation windows. (C) Global arousal and task attention remain intact. Control baseline pupil response, before the embedded pattern and before X (top two panels) and hit rate performance on the X-detection task (bottom) show event rate modulation and dCA1 inhibition do not affect global attention or arousal. **(D)** Maximum pupil size in a 1–2 s window after pattern onset across animals. Boxplots show interquartile range (IQR), whiskers indicate 1.5 × IQR, and red lines mark medians. **(E)** Schematic of optic fibre placement (top). Example histology image showing optic fibre placement (middle). Scale bar, 100 µm. Schematic of optogenetic inhibition experiment (bottom). Light delivered to dCA1 during the exposure phase (100 trials, left) while mice performed the X-detection task. Normal (black) and deviant (red) trials were interleaved. **(F) dCA1 silencing abolishes sensitivity to sequence identity.** Top: PDR to normal and deviant sequences in control sessions (left) and during CA1 inhibition (right). Lines and shaded areas show mean ± SEM across sessions. Bottom: similar to top panels, but in response to X. **(G)** Similar to (C), control measures showing overall attention, and arousal are not affected by task condition and hippocampal inactivation **(H)** Pupil size difference between responses to deviant and normal trials for control (solid) and exposure block inhibition (dashed) sessions. Pupil difference calculated as the difference between the mean response to deviant and normal pattern onset for each session. Lines show mean pupil difference across sessions. Shaded areas show ± SEM. Gray shading marks tone presentation windows. Red shading marks deviant tone presentation window.

In rare versus frequent sessions, bilateral muscimol infusion into dCA1 (Fig. 2A, Fig. S7A) selectively impaired sensitivity to distributional statistics. Compared with control sessions in the same animals, PDRs to frequent sequences increased, abolishing the difference between rare and frequent conditions (shuffle-null permutation test: control, rare > frequent: p = < 1×10⁻⁴; muscimol, rare > frequent: p = 1.0. 10,000 iterations) (Fig. 2B, Fig. 2D). Critically, baseline pupil size, pupil responses to the task-relevant target X, reaction times, and hit rates were unchanged by muscimol infusion (Fig. 2B-C), indicating that dCA1 inactivation disrupted learning of statistical structure rather than altering arousal, attention, or sensory detection.

To dissect the temporal locus of this requirement, we next optogenetically silenced dCA1 (via stimulation of vGAT interneurons) selectively during the learning phase of the normal–deviant paradigm. Light was delivered from −0.25 s to 3 s relative to pattern onset during the initial exposure block (first 100 trials), when animals experienced only the normal ABCD sequence, while dCA1 activity was left intact during the subsequent test block in which normal (ABCD) and deviant (ABAD) sequences were interleaved (Fig.2D). In control sessions, deviants evoked larger PDRs than normal sequences (shuffle-null permutation test: deviant > normal: p < 1×10⁻⁴; 10,000 iterations), consistent with violation of learned expectations. In contrast, when dCA1 was silenced during the exposure phase, this enhanced PDR to deviants was abolished (exposure-block inactivation: deviant > normal: p = 1.0; 10,000 iterations) (Fig. 2E), indicating that transient disruption of dCA1 during learning prevented the formation of the internal model required for subsequent deviance detection. Importantly, optogenetic silencing of dCA1 did not alter baseline pupil size, pupil responses to the X stimulus, reaction times, or hit rates, mirroring the muscimol results (Fig. S2F-G). Thus, dCA1 disruption selectively impaired learning and model formation on long timescales (several trials), rather than producing a general change in arousal or vigilance.

### dCA1 population activity encodes evolving statistical structure

To identify the underlying neural representations supporting SL, we next recorded high-density population activity from dCA1 while mice performed the task (Fig. 3A, Fig. S7B). Individual neurons responded to both the task-relevant X stimulus and the task-irrelevant pattern sequences, though population responses were stronger for X (Fig. 3B). Decoding analyses showed robust stimulus representations (Fig. 3C, Fig. S4B): across all sessions, X could be distinguished from the first pip of the sequence (A) within the first 100 ms (post-sound onset) and, similarly, the first pattern pip could be reliably discriminated from baseline tones (A vs X decoding accuracy, paired t-test: t(520) = 45.53, p = 1.70×10⁻¹⁸²; A vs baseline-tone decoding accuracy, paired t-test: t(520) = 12.79, p = 1.08×10⁻³²). The identity of individual tones within the sequence (A, B, C, D) could also be decoded (4-way tone identity decoding, data vs shuffle, paired t-test: t(520) = 21.05, p = 1.34×10⁻⁷¹) (Fig. 3D, Fig. S4C). Over time, the similarity between different tones within the sequence increased, both within a session and across days (cosine similarity A→B, session 1 vs session 5, paired t-test: t(4) = 4.56, p = 0.0052), suggesting experience-dependent alignment of neural codes for elements belonging to the same statistical structure (Fig. 3E, Fig. S5A).

**Figure 3.**
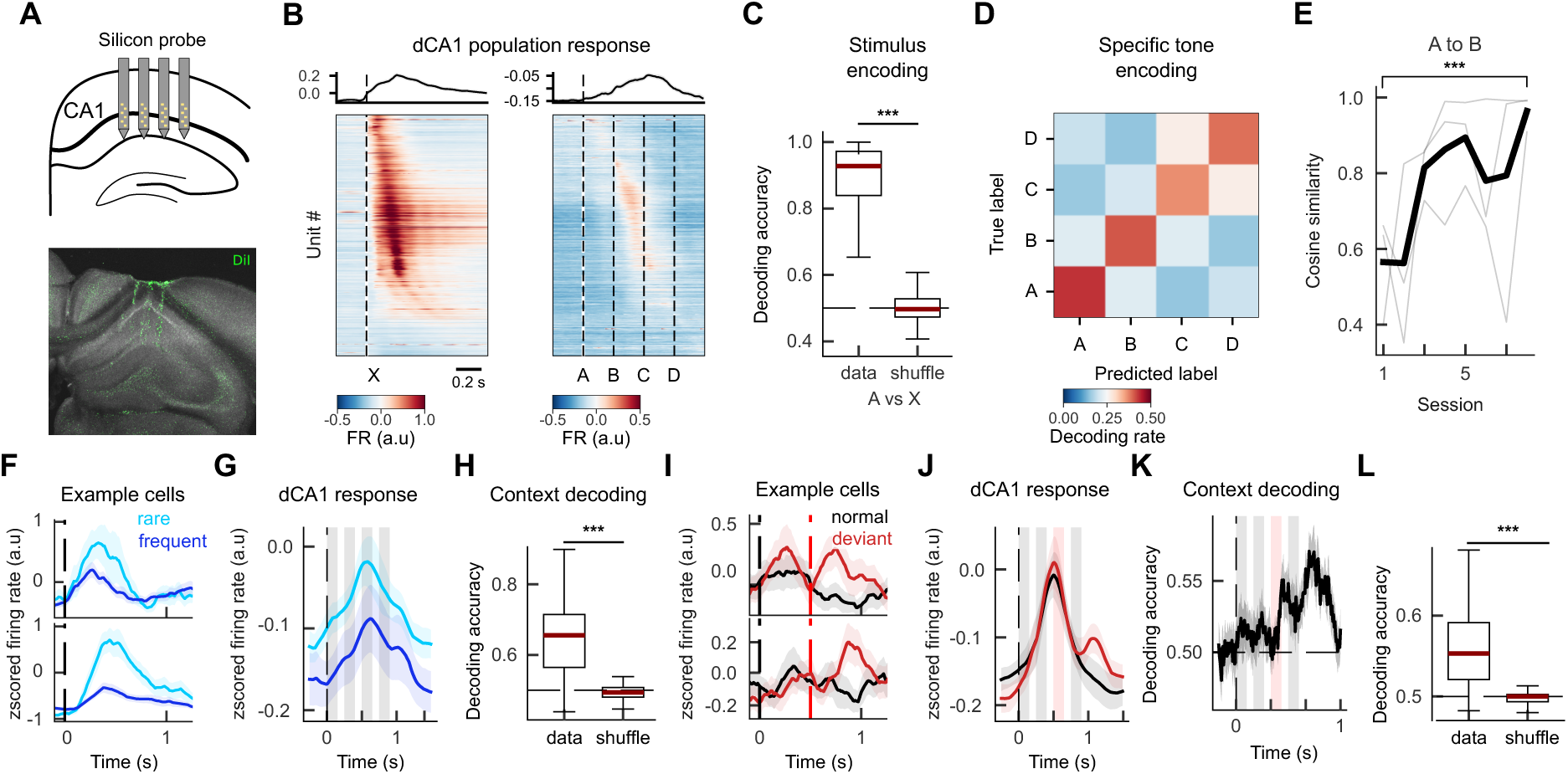
Dorsal CA1 population responses encode both sensory identity and statistical context. **(A)** Schematic of bilateral 4-shank silicon probe recordings from dorsal CA1 (dCA1) and example histology showing DiI-labelled probe tracks. (**B) Sensory responsiveness.** Cross-validated population PSTHs for all recorded units, sorted on odd trials and plotted on even trials. Left, responses to tone X. Right, responses to tones A–D. Top traces show trial-averaged population activity. Vertical dashed lines mark tone onsets. **(C) Sensory decoding.** Decoding accuracy across sessions (n = 343 from 6 mice). Left, binary classification of A versus X using the first 150 ms after tone onset. Right, A versus baseline pip. Boxplots show IQR, whiskers indicate 1.5 × IQR, red lines mark medians, and grey dashed lines indicate chance. Data are compared to label-shuffled controls. ***P < 0.001 (permutation tests). **(D) Tone discrimination.** Confusion matrix for four-way classification of tones A–D using the first 150 ms after tone onset, averaged across sessions. **(E) Stability of sensory representations.** Cosine similarity between population responses to A and B tones across the first five sessions. Population vectors: mean activity 0–0.25 s post-tone onset, averaged across trials. Bold line, mean across mice (n = 3), thin lines, individual mice. **(F-H) Encoding of event rate. (F)** Example z-scored firing rates from two simultaneously recorded neurons aligned to pattern onset, showing their responses modulated by background statistics (rare vs frequent event rate) **(G)** Mean z-scored population activity for A presented in rare (dark blue) or frequent (light blue) contexts (n = 6 mice). Shaded regions indicate SEM; grey bars mark tone presentation windows; vertical dashed line indicates pattern onset. **(H)** Session-wise decoding accuracy (n = 261 sessions from 6 mice) for rare versus frequent contexts compared with shuffled controls using 0 to 1 s post pattern onset window of neural activity. **(I-K) Encoding of sequence identity. (I)** Example z-scored firing rates from two simultaneously recorded neurons aligned to pattern onset, showing their differential response to normal (red) vs deviant (red) patterns. **(J)** Mean population activity for normal (ABCD, black) versus deviant (ABAD, red) sequences. Shading indicates SEM; grey and vertical bars mark tone presentation windows. **(K)** Time-resolved decoding of normal versus deviant sequences using a sliding 250 ms window. Line shows mean decoding accuracy across sessions, shading indicates SEM. **(L)** Session-wise decoding accuracy (n = 23 sessions from 4 mice) for normal versus deviant compared with shuffled controls using 0.5 to 1 s post pattern onset window of neural activity. ***P < 0.001.

Beyond sensory features, dCA1 activity encoded statistical context. Individual neurons modulated their activity (Fig. 3F), and on average neurons increased firing in rare compared with frequent contexts (Fig. 3G), and population activity showed reliable decoding of event probability (rare vs frequent decoding accuracy, data vs shuffle, paired t-test: t(126) = 9.09, p = 1.76×10⁻¹⁵) (Fig. 3H). In the deviant paradigm, elevated responses emerged only after the violating tone (Fig. 3I, Fig. 3I-J). Furthermore, from dCA1 population activity, deviant context could be reliably decoded against normal once the violation had occurred at pip 3 (Fig. 3K, normal vs deviant decoding accuracy over time; Fig.3K data vs shuffle, paired t-test: t(50) = 3.34, p = 1.6×10⁻³). These results show that CA1 simultaneously encodes sensory features and statistical context.

We next asked whether these representations remain stable and adjust flexibly when statistical structure changes without explicit cues (Fig. 4). To test the stability of sensory representations, we trained a decoder to classify individual tones (A, B, C, D) within pattern sequences from dCA1 population activity in the rare context and evaluated its performance in the frequent context. Decoding accuracy remained high, confirming that sensory representations were largely preserved across statistical contexts (Fig. 4A). At the same time, statistical context itself was also reliably decodable from population activity (Fig. 3H), indicating that dCA1 simultaneously represents stimulus identity and the broader statistical context in which stimuli occur.

**Figure 4.**
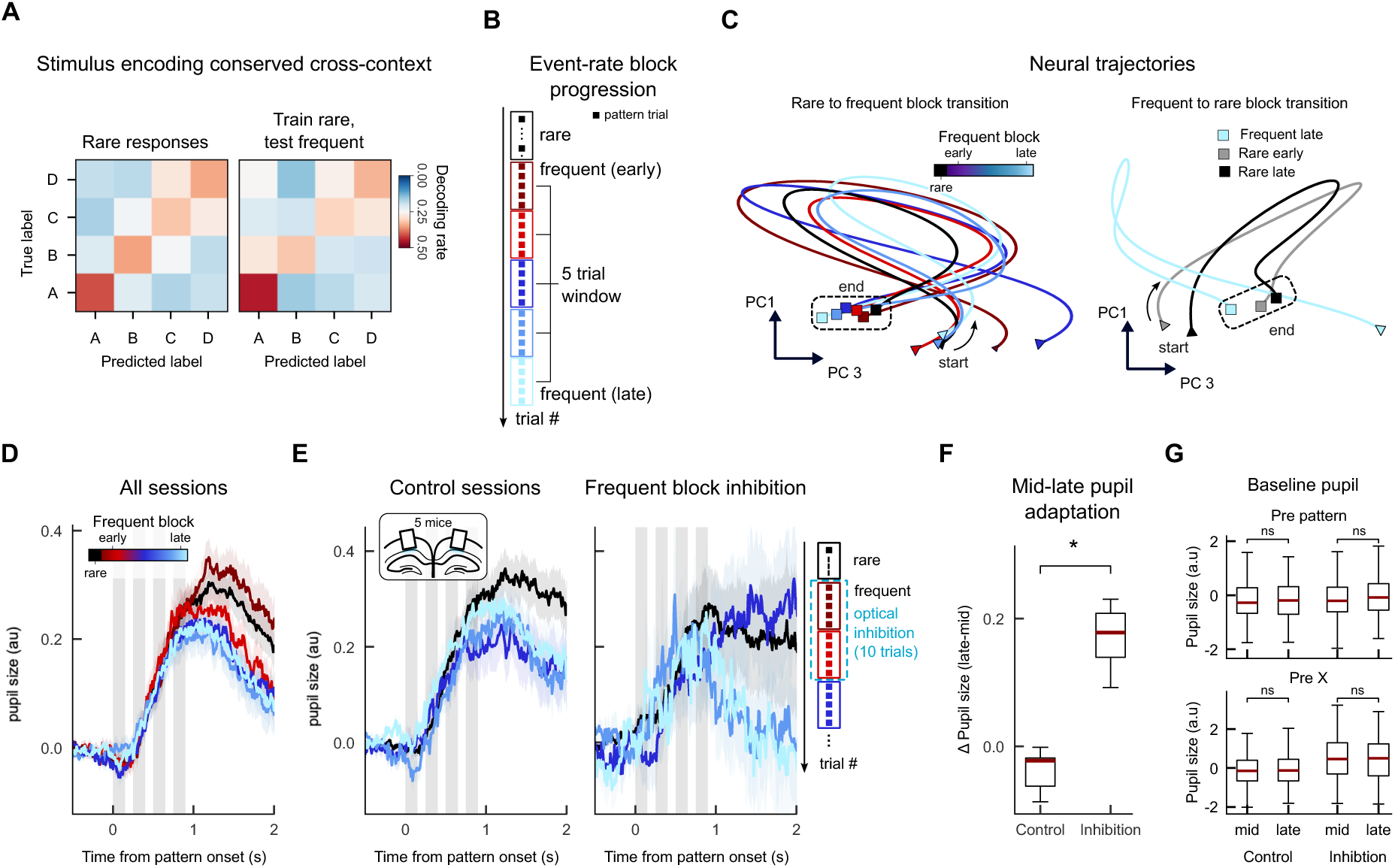
Dorsal CA1 updates internal models via dynamic state-space reorganisation. **(A) Stable sensory coding across statistical contexts**. Confusion matrices for four-way decoding of tones A–D, averaged across sessions. Left, trained and tested using neural responses from the rare context. Right, trained on rare and tested on frequent trials. **(B)** Schematic of event-rate manipulation showing rare and frequent block progression. Frequent block trials grouped in windows of five trials for analysis. **(C) Dynamic reorganisation of the latent state.** dCA1 population responses projected onto the first three principal components. Left: trajectories shown for transition from rare to frequent trials. Right: trajectories shown for transition in reverse, from frequent to rare trials **(D)** Mean pupil responses to frequent patterns grouped by block progression across sessions (n = 10). **E) dCA1 inhibition prevents model updating.** Pupil responses in optogenetic sessions, for rare block (black) and frequent mini-blocks (shades of blue) after termination of opto-inhibition. Left, control sessions. Right, inhibition applied during the first 10 pattern trials of the frequent block. **(F) Quantification of update failure.** Boxplot of the change in pupil size from mid to late frequent trials for control and inhibition sessions. Δ pupil size: difference in mean pupil response (0.5–1.5 s post-pattern onset) between rare and frequent trials, averaged across mice (n = 5). Boxplots show IQR, whiskers indicate 1.5 × IQR, red lines mark medians. **(G) Controls for arousal.** Control baseline pupil responses (before the pattern onset, top; and before the X presentation, bottom) suggest global attention/arousal remains constant under different experimental conditions.

Principal component analysis revealed structured population trajectories evoked by the sequence that diverged between rare and frequent contexts while maintaining a similar overall shape (Fig. 4C-D). Upon uncued transition from rare to frequent blocks (Fig. 4B), trajectories, while following a similar overall shape, gradually shifted along a principal axis of population activity. This emergent dimension (PC3) selectively encoded later portions of the sequence, particularly the final tone (pip 4), and evolved progressively across trials as mice accumulated evidence about the new event statistics (Fig. 4C). PC3 values increased monotonically following the transition, plateauing once the new statistics were established, consistent with the formation of a stable updated internal model (Fig. 4C). A similar pattern was observed following uncued transitions from frequent to rare blocks, where population trajectories shifted in the opposite direction along the same dimension (Fig. 4D), indicating bidirectional updating of statistical representations. These changes in population dynamics closely mirrored changes in pupil responses over time, which were largest immediately after the transition and rapidly decreased and stabilised within approximately ten trials (Fig. 4E). Together, these results show that while core sensory representations remain stable, dCA1 population activity flexibly recruits an additional low-dimensional axis to encode updated statistical structure as experience unfolds, supporting continuous belief updating based on ongoing sensory experience.

To assess whether dCA1 is required for updating internal models in response to changes in statistical structure, we transiently silenced dCA1 optogenetically only during the early portion of the frequent block, immediately following an unmarked transition from rare sequences. Importantly, dCA1 was left intact during the subsequent trials, allowing us to assess the consequences of temporally specific disruption on learning. If dCA1 is necessary for updating beliefs about statistical context, then animals exposed to frequent sequences under dCA1 inactivation should fail to incorporate the new statistics into their internal model. Consistent with this prediction, pupil dilation responses remained elevated on the first trials following the termination of inhibition and were significantly larger than in matched control sessions (Fig. 4E). This indicates that animals failed to register the statistical transition during the inactivation window, despite unaltered sensory input (as indicated by intact behavioural performance on the cover task, Figure 3). To quantify this disruption in learning, we compared average pupil responses across the mid and late phases of the frequent block. In control sessions, pupil dilation was already reduced by the mid-block and remained stable thereafter, indicating that expectations had been updated early following the transition. In contrast, in optogenetic inhibition sessions, pupil responses during the mid-block remained elevated and only significantly declined from mid to late trials (paired t-test, mid-block vs late-block: t(8) = 6.52, p = 0.001) (Fig.4F), indicating delayed belief updating that began only after dCA1 activity was restored. Importantly, baseline pupil size did not differ between control and optogenetic inhibition sessions at any point during the block (Fig. 4G), ruling out changes in global arousal or attentional state as an explanation for the delayed updating. Together, these results demonstrate that dCA1 activity is specifically required during exposure to new statistics to update internal models, rather than for maintaining general arousal, task engagement, or sensory responsiveness.

### dCA1 encodes abstract sequence structure in a factorised population geometry

We next asked whether dCA1 supports not only statistical learning of event probabilities and specific pattern identities, but also abstraction of higher-order structure that generalises across sensory exemplars. Behaviourally, mice showed larger pupil responses to rule-violating than rule-conforming deviants (Fig. 1G), indicating that they had learned an abstract sequence rule (e.g. ‘rising tone pattern’) rather than memorising specific tone transitions.

### Abstraction emerges when the rule becomes inferable

To determine whether dCA1 activity reflected this abstraction, we first analysed population responses in the normal-deviant paradigm (Fig. 5A). Deviants were matched in novelty and pitch shift, dissociating sensory change from rule change. We quantified cosine distance between responses to the learned normal sequence (ABCD₀) and those evoked by rule-conforming (ABCD_1_) or rule-violating (ABBA₁) deviants as the sequence unfolded (Fig. 5C). During the first two tones, when the abstract rule was not yet inferable, population activity was dominated by physical tone properties (Fig.5B), and distances between the normal and both deviants did not differ. By the third tone, when the rule could first be inferred, responses to the rule-conforming deviant became significantly closer to the learned normal than responses to the rule-violating deviant (paired t-tests: ABCD_1_ → ABCD₀ vs ABBA₁→ ABCD₀ at deviant pip 1: t(8)= 1.39, p=0.20; ABCD_1_ → ABCD₀ vs ABBA₁→ ABCD₀ at deviant pip 2: t(8)= 1.54, p= 0.16; ABCD_1_ → ABCD₀ vs ABBA₁→ ABCD₀ at deviant pip 3: t(8)= 10.59, p<0.001; ABCD_1_ → ABCD₀ vs ABBA₁→ ABCD₀ at deviant pip 4: t(8)= 25.38, p<0.001; ABCD_1_ → ABCD₀ vs ABBA₁→ ABCD₀ post deviant pip 4: t(8)= 3.6, p=0.007) (Fig. 5C). This divergence increased as the sequence unfolded, indicating a transition from encoding sensory similarity to encoding inferred latent structure.

**Figure 5.**
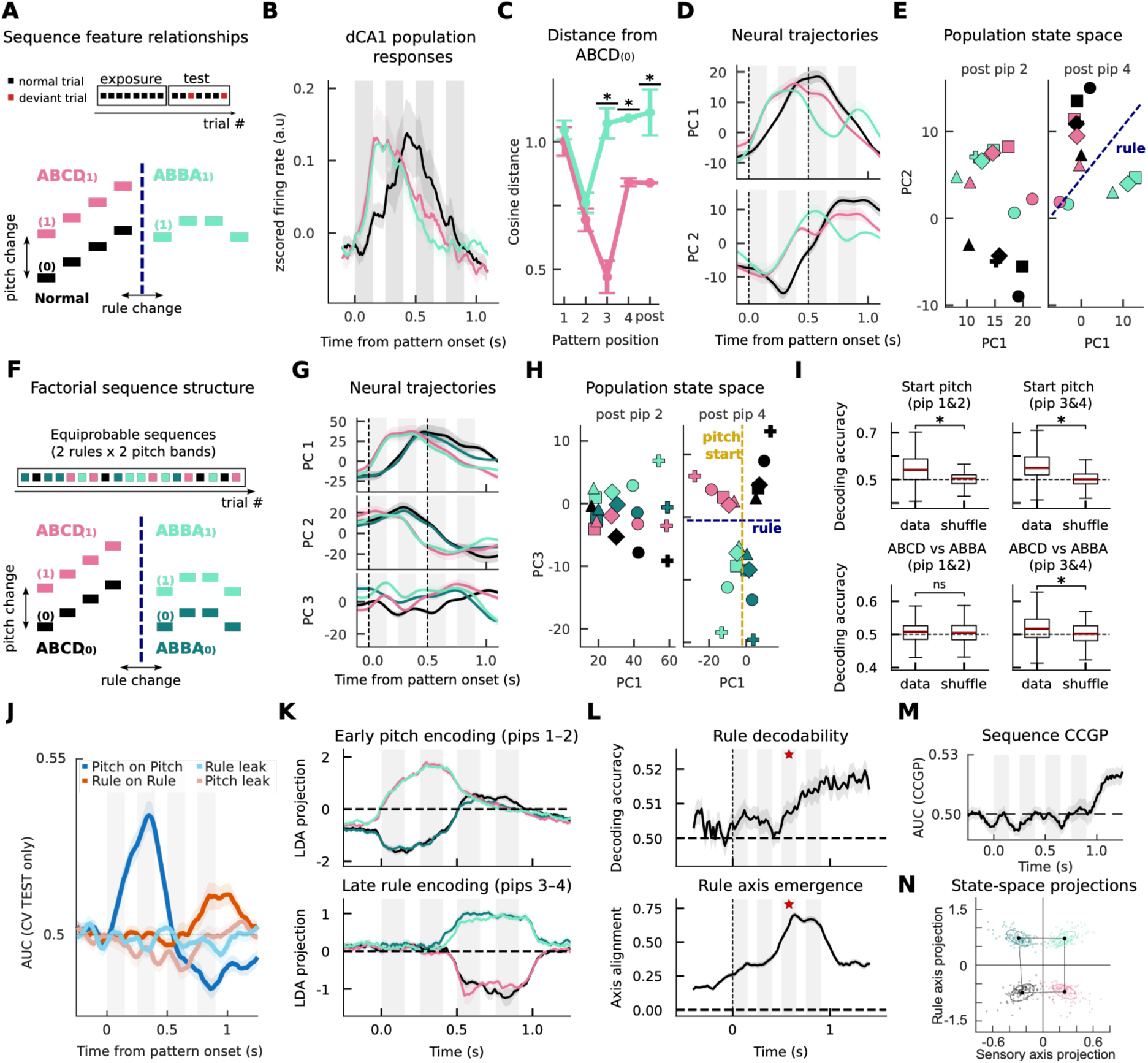
Dorsal CA1 representations transition from sensory feature to rule-based encoding. **(A) Schematic of the generalisation test**. Following exposure to a single normal sequence (ABCD0), two deviant sequences are introduced during the test phase: Rule-Conforming (ABCD1, preserving the rising structure) and Rule-Violating (ABBA1, disrupting structure but matching sensory novelty). **(B) Population response.** Mean z-scored firing rates aligned to pattern onset for normal and deviant sequences. Shaded regions indicate SEM across mice; grey bars denote tone presentation windows. **(C) Distance from Normal pattern.** Cosine distance in principal component space at each tone position, comparing population responses (mean over 100 – 250 ms post tone onset window) for deviant ABCD1 (pink) and ABBA1 (teal) to the normal ABCD0. Error bars show SEM across mice; asterisks denote significant differences. **(D) Population activity projected onto the first two principal components (PC1–PC2)**. PCA was performed separately for each animal (pseudo-population) and trajectories were aligned across animals using orthogonal Procrustes rotation. Shaded regions indicate SEM across mice **(E) State-space snapshots.** Population responses projected into PC space after pip 2 (left) and after pip 4 (right). Colours indicate sequence identity; symbols denote individual animals. **(F) The equiprobable 2x2 design.** Schematic of the factorial design in which four sequences (ABCD0, ABCD1, ABBA0, ABBA1) are presented equiprobably across the session, with two starting pitches and two rules. **(G) Population trajectories (PC1–PC3) for the four equiprobable conditions.** PCs were computed per animal and aligned across animals using orthogonal Procrustes rotation. Shaded regions indicate SEM across mice. **(H) State-space snapshots.** Population responses projected into PC space after pip 2 (left) and after pip 4 (right). Colours indicate sequence identity; symbols denote individual animals. **(I)** LDA decoding accuracy for pitch (top) and rule (bottom) during early (pips 1–2) and late (pips 3–4) stimulus periods. Classifiers were trained on population responses (0-500 ms for pip1/2; 500-1000 ms post pattern onset for pip 3/4) and evaluated using 5 -fold cross-validation. Box plots show decoding performance across sessions; shuffle controls indicate chance-level decoding. Red lines mark the median. Boxes indicate interquartile range, whiskers denote 1.5 x IQR across sessions. **(J–K) Orthogonal Factorisation. (J) Time-resolved decoding on LDA-defined axes**. Discriminability was quantified at each time point as AUS from test-set projection scores. Four curves were computed per session: Pitch on Pitch axis (low vs high on pitch axis, trained on pip1-2, dark blue), Rule on Rule axis (ABCD vs ABBA on rule axis, trained on pip 3-4, dark red), and two cross-axis controls (Pitch on Rule axis, light red; and Rule on Pitch axis, light blue) that report “leakage”, which remains near chance throughout. Lines show mean +/- SEM across sessions. **(K) Population responses projected onto linear discriminant (LDA) axes**. Top: Projection onto the pitch LDA axis trained on responses during pips 1–2. Bottom: Projection onto the rule LDA axis trained on responses during pips 3–4. Traces show mean across sessions; shaded regions indicate SEM. Gray bars mark tone presentation windows. **(L) Rule axis emergence.** Top: Cross-validated LDA decoding accuracy for rule identity computed in sliding time windows (chance = 0.5, dashed line). Bottom: Rule axis emergence, quantified as the cosine alignment between LDA axes trained at each time window (sliding 200 ms window) and a late reference rule axis (trained on pips 3–4). Star marks discoverability of rule at pip 3. **(M) Cross-condition generalisation performance** (CCGP; AUC) for rule identity over time, training the classifier within one pitch band (ABCD0 vs ABBA0) and testing on the other (ABCD1 vs ABBA1), and vice versa (then averaged); curves show mean ± SEM across sessions **(N) The geometry of abstraction.** Projection of pip 1-2 on Pitch axis, and pip 3-4 on Rule axis. Dots show individual sessions (n = 136, train data).

Principal component projections confirmed this temporal reorganisation (Fig. 5D-E). Early in the sequence, projections onto the first two principal components grouped sequences primarily by pitch: responses to ABCD₁ and ABBA₁ overlapped and were distinct from ABCD₀ (Fig. 5D). After pip 3-4, this geometry reorganised such that responses to ABCD₀ and ABCD₁ converged, while ABBA₁ separated, consistent with rule-based grouping (Fig. 5D, right). Across animals, pooled PC1-PC2 projections showed clustering by pitch early (post–pip 2, before pip 3) and by abstract rule late (post-pip4, Fig. 5E). Thus, abstraction appears precisely when it becomes computationally inferable.

### Rule coding is not driven by deviance or surprise

To directly test abstraction and generalisation independently of novelty, deviance or expectation violation, we introduced a fully equiprobable 2 x 2 design (Fig 5F). Two abstract rules (ABCD and ABBA) were each instantiated by two pitch exemplars (0 and 1), yielding four equally probable sequences (ABCD₀, ABCD₁, ABBA₀, ABBA₁). Because all four sequences were presented with equal probability, any separation in neural state space could not be attributed to surprise, rarity or deviance detection. Similar to the previous normal-deviant paradigm, during the first two tones, when the abstract rule was not yet inferable, population activity was dominated by physical tone properties (Fig. S6A). Time-resolved projections into the first three principal components revealed dissociable sensory and abstract dimensions (Fig. 5G). During the first two tones (pip 1-2), PC1 (and PC2) encoded pitch. After pip-3, when rule identity becomes inferable, PC1 and PC2 continued to encode the initial pitch band (defined by pip1-2), preserving the starting sensory context across the unfolding sequence. In contrast, PC3 showed little structure early on but, following pip 3, selectively separated sequences by abstract rule irrespective of pitch. In PC1-PC3 space (Fig. 5H), late population responses occupied four distinct quadrants, with the horizontal axis (PC1) encoding pitch band and the vertical axis (PC3) encoding rule identity (Fig. 5J). This late-emerging rule separation in an equiprobable design demonstrates that dCA1 encodes abstract structure independent of novelty or expectation violation.

### Sensory and rule axes are dissociable and orthogonal

To explicitly test separability of sensory and abstract codes, we performed cross-validated discriminant analysis (LDA) on population responses in the equiprobable 2x2 design (Fig. 5I-K). Decoders were trained separately to classify pitch band (0 vs 1, pooled across rules) and abstract rule (ABCD vs ABBA, pooled across pitch), in early (pips 1-2) and late (pip 3-4) windows. During the early window, pitch decoding was robustly above chance, whereas rule decoding was at chance (early pitch decoding: t(270)=7.24, p<0.001, early rule decoding: t(270)=0.06, p=0.52). During the late window, after the third tone revealed the sequence rule, rule decoding rose significantly above chance while pitch decoding remained high (late pitch decoding: t(270)=9.55, p<0.001, late rule decoding: t(270)=3.74, p<0.001) (Fig. 5I, see Fig S6B for pip decoding over time). Thus, sensory information is available from the outset whereas abstract rule information emerges only once sufficient sequential evidence accumulates. Importantly, the pitch and rule axes were approximately orthogonal, with minimal leakage: decoding of rule along the pitch axis and pitch along the rule axis remained near chance (Fig. 5J). Population projections onto these axes showed early separation by pitch and late separation by rule (Fig. 5K), consistent with a factorised representational geometry.

We next quantified how rule information evolves over time using sliding-window cross-validated LDA (Fig. 5L, top). Rule decoding remained at chance during pips 1-2, rose sharply after pip 3, and continued to increase toward the end of the sequence. Rule information therefore emerges online as evidence accumulates. To test whether this reflects formation of a stable low-dimensional axis, we defined a reference rule axis from the late window (pip 3-4) and computed cosine alignment between axes trained at each time point and this reference (Fig. 5L, bottom). Alignment was low early, then progressively increased and peaked at sequence end. Notably, the rise in axis alignment closely tracked the increase in decoding accuracy, indicating that improved rule decodability reflects stabilisation of a consistent low-dimensional rule axis (see Fig. S6C for pitch axis emergence).

### Rule representations generalise across pitch exemplars

Finally, we quantified cross-condition generalisation performance (CCGP) (Bernardi et al., 2020). A classifier trained to distinguish rule identity within one pitch band (ABCD₀ vs ABBA₀) was tested on the other (ABCD₁ vs ABBA₁), and vice versa (Fig. 5M). Rule generalisation rose selectively after pip 3–4, demonstrating that the neural code for rule transfers across sensory instantiations. Mean AUC in the late window (0.80–1.20 s) was 0.5097 ± 0.0033 (SEM; n=135 sessions), significantly above chance (t(134)=2.99, p=0.0034) and confirmed by label-shuffle permutation testing. This analysis directly establishes that the late rule representation is shared across pitch instantiations, a hallmark of abstraction that cannot be demonstrated by within-condition decoding alone.

Finally, we examined the geometry of the rule and pitch representations. In the two-axis state space defined by the pitch and rule discriminant axes, the four conditions formed an approximately rectangular arrangement late in the sequence (Fig. 5N): low-pitch conditions occupied the left, high-pitch conditions the right, while ABCD and ABBA were separated along an orthogonal rule dimension. The pitch difference vector (ABCD₁−ABCD₀) was approximately parallel to the pitch difference vector (ABBA₁−ABBA₀), consistent with a factorised code in which rule acts as a translation that preserves pitch structure across contexts (parallelism/rectangle metrics quantified across sessions, Fig S6D). Together, these convergent analyses show that dCA1 population activity preserves stimulus identity while constructing an orthogonal representation of latent sequence structure that emerges online as evidence accumulates, enabling generalisation across sensory exemplars without requiring novelty or deviance.

Together, these analyses show that dCA1 does not merely signal novelty or deviance. Instead, hippocampal population activity reorganises dynamically to encode inferred latent structure in a distinct low-dimensional subspace that emerges when sufficient evidence accumulates. Sensory and abstract variables are encoded along dissociable, approximately orthogonal axes, enabling generalisation across exemplars while preserving stimulus identity. This factorised geometry provides direct neural evidence that the hippocampus constructs abstract internal models of environmental structure.

## Discussion

A fundamental challenge in systems neuroscience is to understand how the brain extracts structure from experience when learning is unguided by goals, feedback, or reinforcement. Here, we demonstrate that dorsal CA1 (dCA1) plays a causal and mechanistic role in this process, supporting the formation, updating, and generalisation of internal models of auditory regularities under fully unsupervised conditions. By combining a reinforcement-free behavioural paradigm with temporally precise optogenetic perturbations and large-scale recordings, we identify how hippocampal population dynamics implement statistical learning beyond simple novelty detection.

Our results uncover multiple levels of hippocampal involvement in statistical learning, from the acquisition of new regularities to the abstraction of shared structure across experiences. During acquisition, dCA1 activity was required for learning new statistical regularities: optogenetic inactivation during early exposure to a novel context abolished later learning-related pupil responses, indicating that belief updating was prevented when dCA1 activity was suppressed. Crucially, task accuracy, reaction time, baseline pupil size and pupil responses to the task-relevant target remained intact. These controls rule out nonspecific effects on global arousal, attention or task engagement, demonstrating that dCA1 is selectively required for building and updating internal models of environmental structure.

At the neural level, dCA1 population dynamics jointly encoded tone identity and statistical context. Sensory representations of individual tones were stable across contexts, yet population trajectories diverged systematically depending on the underlying statistics. Crucially, changes in statistical structure were not reflected as loss of sensory coding, but as structured reorganisation within population state space.

A unifying feature across our manipulations is that learning new statistics was accompanied by the emergence of new, low-dimensional population axes that encoded statistical context on top of preserved sensory representations. In the rare-frequent manipulation, uncued changes in event rate led to gradual shifts in population activity along a distinct principal component, mirroring the time course of behavioural updating. Similarly, in the abstraction experiments, abstract sequence structure (ABCD vs ABBA) was captured along a separate population dimension that emerged only once the rule became inferable, while sensory features such as pitch remained encoded in orthogonal dimensions. Despite probing different forms of structure (event probability versus abstract generative rules) both manipulations revealed the same organising principle: dCA1 does not overwrite sensory codes when statistics change but dynamically expands its representational space to incorporate new contextual dimensions.

This factorised population geometry provides a mechanistic account of abstraction and generalisation. By encoding sensory features and latent structure in dissociable subspaces, dCA1 can generalise across physically distinct exemplars that share the same underlying rule, while preserving information about stimulus identity. This was directly demonstrated using an equiprobable 2×2 design, where all sequences were likely and no deviance or surprise-based explanation applied. In this setting, sensory pitch and abstract rule were represented in dissociable, approximately orthogonal subspaces. A late-emerging rule axis generalised across pitch exemplars, as quantified by cross-condition generalisation performance (CCGP), demonstrating that the hippocampal code captured latent structure rather than stimulus-specific features or deviance. Together, these findings indicate that dCA1 supports abstraction by dynamically expanding its representational geometry into factorised subspaces, enabling generalisation without overwriting sensory codes. These hierarchical operations occurred without task relevance, feedback, or reward.

Converging evidence has implicated the hippocampus in predictive and latent-state learning across diverse behavioural domains. Our study isolates this process in its canonical form—incidental, reinforcement-free exposure to structured input—revealing hippocampal mechanisms that support structure learning independent of task demands. These findings complement work on latent state learning (Mishchanchuk et al., 2024; Sun et al., 2025) and predictive modelling in hippocampus (Miller et al., 2023) and support the theoretical accounts in which hippocampal circuits compress regularities across experience into flexible representations for inference (Behrens et al., 2018; Buzsáki & Tingley, 2018; Chen et al., 2024; Kanter et al., 2025; Levenstein et al., 2024; Stachenfeld et al., 2017).

Importantly, our results go beyond framing dCA1 as a novelty or mismatches detector (Knight, 1996; Kumaran & Maguire, 2006, 2009; Vinogradova, 2001; Yi et al., 2022). Although this mismatch-based framing has been influential, it does not explain how expectations are formed, integrated across discontinuous experiences, or generalised across abstract structure. Neural correlates of abstract task structure and latent states in dCA1 (Jeong et al., 2018; Nieh et al., 2021; Sun et al., 2025) instead point to a more expansive function as a hub for learning. Within this framework, our finding that disrupting dCA1 during initial exposure prevents subsequent updating demonstrates that incorporating new regularities depends on intact dCA1 function. This extends observations from spatial and decision-making domains to statistical learning and underscores the hippocampus’s capacity to integrate related experiences through shared latent structure.

Our results further suggest that CA1 may function as a general-purpose statistical learning system whose population geometry continuously adjust to reflect ongoing environmental statistics. The gradual drift in CA1 population structure across block transitions paralleled behavioural updating of expectations and resembles the representational drift frequently reported in hippocampal recordings (Geva et al., 2023; Kentros et al., 2004; Ziv et al., 2013). Rather than reflecting noise or instability, such drift may index principled belief updating as internal models are refined over time.

At the circuit level, these findings provide an empirical basis to revisit long-standing theoretical distinctions within the hippocampal network (McClelland et al., 1995). The trisynaptic DG–CA3–CA1 loop is classically associated with episodic memory and pattern completion, whereas the monosynaptic entorhinal–CA1 pathway has been proposed to support online comparison between sensory input and internal predictions (Koster et al., 2018; Schapiro et al., 2017). Our data are consistent with aspects of this division: dCA1 activity reflected both sensory identity and evolving context, and optogenetic inactivation during context transitions disrupted belief updating. Notably, the enhanced pupil responses we observe to rare or structurally violating events resemble prediction-error signals widely reported in sensory and motor cortices (Audette et al., 2022; Fiser et al., 2016; Furutachi et al., 2024; Keller & Mrsic-Flogel, 2018), suggesting that hippocampal computations may interface with distributed cortical mechanisms for updating internal models when expectations are violated (Yi et al., 2022). Together, these results motivate further investigation into how hippocampal, entorhinal and sensory circuits interact to extract and infer structure from experience.

Finally, our study advances the investigation of statistical learning beyond the constraints of existing paradigms. Many rodent tasks designed to probe structure learning rely on reinforcement, feedback, or task relevance (Fortin et al., 2002; Menichini et al., 2025; Shahbaba et al., 2022), making it difficult to disentangle structure learning from reward-driven updating or performance monitoring. While predictive-coding and oddball frameworks reveal sensitivity to local or global deviations (Meyer & Olson, 2011; Murphy et al., 2008; Toro et al., 2005), they typically operate over short temporal windows, and often lack a direct measure of learned knowledge. In contrast, the regularities in our paradigm (i.e. event rate and abstract rule) can only be inferred across multiple sequences, separated by intervening events. Learning therefore depends on integrating information over extended timescales, rather than detecting isolated sensory changes. Our reinforcement-free, task-irrelevant design captures how organisms extract such latent structure without instruction or feedback, and continuous physiological readouts—such as pupil-linked surprise—allow us to track the gradual formation and updating of internal models without requiring any behavioural report. Together, these features enable access to forms of statistical learning that extend beyond deviance detection to the acquisition of abstract, higher order latent structure.

Together, these results establish dCA1 as a critical substrate for unsupervised SL in the rodent brain, operating across timescales and levels of structure, and independent of reinforcement. By bridging behavioural, physiological, and circuit-level measurements across species, our study opens a new avenue for investigating how the brain implements flexible, abstract structure learning, without reward, task, or explicit feedback.

## Supplementary Materials – Methods

### Animals

All experiments were approved by the Sainsbury Wellcome Animal Welfare Ethical Review Body and conducted under the UK Animals (Scientific Procedures) Act of 1986 (PPL PE4FA53CB). Male and female C57BL/6J mice (24 mice) (6–12 weeks old, Charles River Laboratories) were used. Animals were housed on a 12-h reverse light/dark cycle, group-housed when possible (2–4 per cage) or single-housed when chronically implanted. Body weight was maintained at approximately 85% with water restriction, with ad libitum access if weight fell below 80%. Mice were habituated to handling and water restriction for at least 48 hours prior to behavioural training.

### Surgical procedures

#### Head-bar implantation

Mice were anesthetized with isoflurane (5% induction, 2% maintenance) and body temperature maintained at 37°C. Pre- and post-operative analgesia (meloxicam, 5 mg/kg subcutaneous) was administered. Scalp was shaved, sterilized, and the skull exposed. The skull was etched, and a stainless-steel head-bar was affixed with dental cement. Animals recovered in heated cages with wet food and were returned to home cages after ambulation.

#### Infusion cannula implantation

Bilateral dorsal CA1 cannulas were implanted at AP −2.1 mm, ML ±1.8 mm, DV −1.2 mm relative to bregma. Craniotomies were drilled with a 0.8 mm spherical bur, etched, and cannulas lowered to target depth. Implants were secured with dental cement, and a head-bar was affixed rostral to the cannula.

#### Optic cannula implantation

Optogenetic experiments were conducted using transgenic mice expressing channel rhodopsin in inhibitory VGAT neurons (VGAT-ChR2-YFP; Jackson Laboratories, USA). Optic cannulas (2mm long,400 µm core, 0.39 NA; Thorlabs) were implanted at AP −2.1 mm, ML ±1.7 mm, DV −1.1 mm relative to bregma. Cannulas were implanted at a 10° angle. Craniotomies (ML position 1.9 mm) were drilled with a 0.8 mm spherical bur, etched, and cannulas lowered to target depth. Implants were secured with dental cement, and a head-bar was affixed rostral to the cannula.

#### Chronic probe implantation

Chronic 4-shank silicon probes were implanted above dorsal CA1 (AP -2.1 mm, ML - 1.5 mm, DV 0.8–0.9 mm below brain surface), coated with DiI. Craniotomies were drilled and probes were lowered at 2–3 μm/s. A ground screw was inserted near bregma. Implants were fixed with UV-curable cement, connectors secured to the head-bar, and wrapped with surgical tape. Animals were single-housed post-surgery. Probes were lowered across multiple days to the CA1 pyramidal layer (estimated DV of 1.4 mm below brain surface). Position of probe sites in CA1 pyramidal region was confirmed by visual inspection of LFP signal.

### Behavioural assays

#### Head-fixed apparatus

Experiments were conducted in a custom soundproof chamber (650 × 600 × 650 mm). Mice were head-fixed on a platform with paw support and placed in front of a lick spout. Licks were detected using capacitive or photo-interrupter sensors. Water rewards (1–10 μL) were delivered via solenoid valves calibrated with a USB scale. Auditory stimuli were delivered via stereo speakers controlled by Harp devices . Pupil videos were acquired using an infrared-illuminated camera. Ambient light was controlled to maintain medium pupil dilation.

#### X- detection task design

Subjects performed a simple auditory target detection task (tone X in a stream of background sounds) while being passively exposed to structured, task-irrelevant tone sequences (e.g., ABCD, rising tones). Trials consisted of a stream of repeated pure tones (base tone: 5275 Hz, 150 ms on / 100 ms off) with a target broadband noise (“X”: 20 Hz–20 kHz, 250 ms) embedded at a random time between 5–10 s after trial onset. On a subset of trials, pattern tone sequences were embedded at random times, non-predictive of X. Correct licks within a 1-s response window post X presentation triggered water reward (1–10 μL) and a 4-pip auditory cue (4000 Hz, 4 × 100 ms pulses with 100 ms gaps). Premature licking delayed X onset.

To maximise the attention paid to all auditory stimuli, catch trials where the pattern sequence was interrupted by the presentation of the X or by returning to the presentation of the baseline tone were randomly interleaved. Catch trials made up ∼10% of all trials. Trials where the pattern was a deviant sequence were never interrupted.

#### Pattern sequences

Embedded tone sequences (normal: ABCD; deviant: ABAD; additional: A_1_B_1_C_1_D_1_, A_1_B_1_B_1_A_1_) were presented independently of X. Sequences consisted of four 150-ms tones separated by 100-ms gaps. Deviant sequences altered the third pip or violated the sequence rule.

#### Training procedure

Animals underwent a multi-stage training protocol designed to shape performance on the final task. The progression criteria for each stage are summarized in Table 1. Mice were first habituated to the head-fixed setup and trained to lick water rewards from the spout. They then entered training stage 0, in which the X stimulus was presented in isolation at random intervals. Early in this stage, each X presentation was automatically paired with a water reward. As training progressed, trials requiring the animal to lick in response to X for reward delivery were gradually introduced, while the proportion of automatic reward trials was correspondingly reduced.

**Table 1:** Training stages for animal shaping for X-detection task.

| Stage | Description | Exit criteria |
| --- | --- | --- |
| -1 | Habituation to head-fixation and water reward | Animal licks spout upon water delivery |
| 0 | Animal licks in response to X presentation in isolation without cueing. Proportion of cued trials ramped down. | Animal correctly licks to X on un-cued trials on at least ~75% trials. Total correct trials ~200 trials. |
| 1 | Auditory stream with repeating baseline tone introduced. Random trial lengths introduced. Licking outside of response window increasingly penalised. | Animal's licking restricted to response window on ~90% of trials. Animal can withhold licking on trial durations up to 12 seconds. |
| 2 | Embedded pattern tones introduced | na |

**Table 2:** Trial design parameters.

| Parameter | Details |
| --- | --- |
| Base tone | 5275 Hz, 150 ms on / 100 ms off |
| Target X | White noise, 20 Hz–20 kHz, 250 ms |
| Reward cue | 4000 Hz, 4 × 100 ms pulses, 400 ms total |
| Normal sequence (ABCD) | 5920, 7902, 10548, 14080 Hz; 4 × 150 ms / 100 ms off |
| Deviant sequence (ABAD) | 5920, 7902, 5920, 7902 Hz; 4 × 150 ms / 100 ms off |
| Pitch shifted normal (ABCD(1)) | 7548, 9956, 13289, 17739; 4 × 150 ms / 100 ms off |
| Pitch shifted deviant (ABBA(1)) | 7548, 9956, 9956, 7548; 4 × 150 ms / 100 ms off |
| Event rate tracking | Interleaved blocks of high (0.9) and low (0.1) pattern presentation rates |
| Normal vs. deviant sequences | Exposure phase with normal sequences (100 trials, rate 0.9), test phase with deviant sequences (rate 0.1–0.2), minimum 5 trials between deviants |
| Rule abstraction | Multiple sequences presented randomly; stimuli shared either generative rule or frequency group; non-pattern trials comprised 20% |
| Passive presentation | Tones from that day's session presented without reward; interstimulus interval randomized 3–5 s; 300 presentations (~20 min) |

After mice reliably licked in response to the X stimulus, they advanced to stage 1, where they performed the cover-up task without the embedded pattern sequence. At the start of this stage, water rewards were again automatically delivered after X presentation, and the rate of automatic rewards was gradually decreased across correct trials as in stage 0. A penalty for licking outside the response window was also introduced, implemented as a delay in trial onset. This penalty was initially 0.5 s and was gradually increased to 2 s over training. Through this stage, animals learned to lick selectively following X presentation and to suppress licking outside the response window.

Once mice showed stable, criterion-level performance on the X stimulus alone, they proceeded to the full task incorporating the embedded pattern sequence. Animals typically completed all training stages within 5–10 sessions.

#### Task configurations

After training on the X-detection task and reaching criterion performance, mice completed several task variants depending on the experiment (Fig. 3C).

#### Event rate tracking

Sessions consisted of interleaved blocks with high (0.9) or low (0.1) pattern presentation rates. Each session began with 10 warm-up trials during which no pattern sequences were presented. Following warm-up, the probability that the embedded pattern sequence preceded the X stimulus was set according to the current rate. Block durations were adjusted so that 10–30 pattern sequences occurred per block. Rare and frequent block types alternated until mice reached satiation (∼300 trials per session).

#### Normal vs. deviant sequences

Sessions comprised two main phases: an exposure phase and a test phase. During exposure (100 trials), mice were presented with a standard (normal) pattern sequence at a constant presentation rate of 0.9.

Two configurations of test phases were used. In one, a single deviant pattern (ABAD) was introduced, differing from the normal sequence (ABCD) by the identity of the third pip. The deviant presentation rate was 0.1, with at least five trials separating consecutive deviants. In the second configuration, the normal sequence (A₀B₀C₀D₀) was paired with two deviant types: a rule-conforming deviant (A₁B₁C₁D₁) and a rule-violating deviant (A₁B₁B₁A₁). Deviants occurred with a rate of 0.2, with one of the two selected at random on each deviant trial. Consecutive deviants were separated by a minimum of five trials.

#### Behavioural Control

Protocols were implemented in Bonsai, which controlled lick detection, water delivery, TTL triggers, and stimulus presentation. Data were logged for offline analysis.

#### Muscimol infusion

Mice were infused with muscimol (1mg/ml) prior to the session. Mice were head-fixed before infusing 0.5 µL into each cannula using a Hamilton syringe. Mice were returned to their cage for 30 minutes before beginning the training session.

#### Optogenetic stimulation

Blue light (470 nm, 4 mW at fibre tip) was delivered bilaterally using the Thorlabs optogenetic system. On trials with optogenetic perturbation, stimulation was triggered 0.25 s pre pattern onset to 3 s post pattern onset followed by a 250 ms ramp down. The laser fluctuated according to a bilateral 40 Hz square pulse pattern with a 50% duty cycle. To prevent the mice seeing the blue light, the fibres and skull area were covered with foil. Trials with optogenetic stimulation were not included in the PDR analysis.

#### Pupil Imaging and Preprocessing

Videos of the left pupil were recorded using a USB camera and an IR lamp for illumination. A visible lamp was also used to control the background luminance. Pupil diameter was measured using PupilSense, a pretrained Detectron2 model (Islam et al., 2024; Wu et al., 2019). Raw pupil traces were uniformly resampled to 90 Hz. Outliers and missing values were removed and interpolated following procedures previously described (Leys et al., 2013). Traces were z-scored using the mean and standard deviation of each session.

Pupil data were analysed using custom Python scripts. To compare dilation responses across event conditions and trial types, the z-scored pupil trace was epoched from 1 s before to 3 s after each event. Traces were baseline-corrected by subtracting the mean pupil size within the 1 s pre-event window. Epochs were excluded if more than 50% of samples were marked as outliers, or if over 50% of baseline samples were identified as outliers.

#### Quantification of condition-dependent pupil responses

To quantify condition-dependent pupil changes, the mean pupil response was computed across trials within each condition for each session. Differences between conditions were evaluated using a cluster-based permutation test implemented in MNE-Python. Analyses were performed on pupil traces spanning 0–3 s post-event. Observed *t*-values were computed for each time point, and clusters of temporally contiguous samples exceeding a threshold (*t* > 2.5) were identified. For each cluster, the cluster mass was defined as the sum of *t*-values within that cluster. The observed cluster masses were compared against a null distribution generated from 10,000 iterations of trial label permutation. A one-tailed test (positive clusters only) was used, and empirical *p*-values were calculated as the proportion of permutations with cluster mass greater than or equal to the observed value.

### Electrophysiological data analysis

#### Neural recordings and preprocessing

Neural recordings were obtained using dual 4-shank silicone probes (16 recording sites per shank; 250 μm inter-shank spacing; 150 μm spacing between probes; ASSY-236-P1, Cambridge NeuroTech) or double-sided 4-shank probes with identical shank spacing (ASSY-325D, Cambridge NeuroTech). Signals were acquired at 30 kHz using *Open Ephys* DAQ hardware and acquisition software. Preprocessing was performed in *SpikeInterface (Buccino et al., 2020)*, including re-referencing to the common average per shank and bandpass filtering between 300–9000 Hz. Spikes were sorted using *Kilosort4 (Pachitariu et al., 2024)*, and unit waveforms were manually inspected in *SpikeInterface*. Units were excluded if they exhibited abnormal waveforms or had mean firing rates below 0.01 Hz.

#### Firing rate estimation

Peristimulus firing rates were computed in 10 ms bins and smoothed by convolution with an exponential kernel (width = 40 ms). Firing rates were then z-scored relative to a baseline period of at least 2 s preceding stimulus onset. All analyses were performed using custom Python scripts and *scikit-learn (Pedregosa et al., 2011)*.

#### Population decoding

For population decoding, mean firing rates were computed within defined temporal windows (typically 0–250 ms post-stimulus onset, unless otherwise stated). Session-level decoding was implemented using L2-regularised logistic regression (*scikit-learn*). For binary classifications, the model fit a single logistic function to separate the two classes. For multiclass decoding (A, B, C, D), a multinomial (softmax) logistic regression was used, jointly optimising class-specific weight vectors to estimate the probability of each class. Decoding performance was evaluated using five-fold cross-validation. Condition decoding was performed using Monte Carlo cross-validation with 80/20 train–test splits over 1000 iterations. To assess significance, decoding was repeated on label-shuffled data using the same cross-validation procedure, and decoding accuracies were aggregated across sessions. Real and shuffled accuracies were then compared using an independent *t*-test. Chance levels were defined as 50% for binary and 25% for four-class decoding.

#### Principal component analysis (PCA)

For PCA, responses were z-scored within each trial using the trial’s mean and standard deviation. A pseudo-population was constructed for each all animals, and PCA was computed on trial-averaged responses. To compare trajectories across mice, the projected population trajectories were aligned into a common space using a linear alignment fitted within a chosen time window (0 – 1 s post pattern onset using orthogonal Procrustes). This alignment was then applied to the full trajectories. For 2-D visualisation, the first two aligned PCs were plotted, and trajectories were smoothed with a 10-sample (100 ms) Gaussian filter for display only.

For distance analyses, each event was represented by the mean 15-dimensional PC vector computed over the 0–150 ms window after tone onset, separately for conditions A, B, C, and D. Cosine distance was then computed between pairs of these vectors, where 𝑥 and 𝑦 were the mean PC vectors for two conditions, defined as 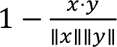, with a minimum and maximum distance of 0 and 2 respectively.

#### Linear discriminant analysis (LDA)

We used linear discriminant analysis (LDA) to identify low-dimensional axes capturing sensory (pitch) and abstract (rule) information in dCA1 population activity, and to track how these axes evolve over the course of a sequence pattern. Analyses focused on an equiprobable 2×2 design with four conditions: rising sequences (A_0_B_0_C_0_D_0_, A₁B₁C₁D₁) and rising–falling sequences (A_0_B_0_B_0_A_0_, A₁B₁B₁A₁), where subscripts 0/1 denote the two pitch exemplars. For each session, we constructed trial-wise population response vectors from the activity matrix (time × units × trials). Unless otherwise stated, activity was baseline-subtracted by subtracting the mean activity in a pre-sequence baseline window from each neuron on each trial. For time-resolved analyses we either used individual time bins or a short sliding integration window, computing the mean activity across the window for each neuron to yield one population vector per trial per time point. Units were z-scored using statistics computed on the training data only (mean and standard deviation across training trials), applying the same normalisation to the held-out test trials. Units with very low training-set variance (standard deviation below a fixed floor of 1e-3) were excluded.

We defined two binary classification problems, each trained in a pre-specified time window:

1. **Pitch axis (sensory axis).** An LDA classifier was trained to discriminate pitch exemplar (low vs high) using trials pooled across both rules. The pitch axis was fit using population activity in an early window spanning pips 1–2.
2. **Rule axis (abstract axis).** An LDA classifier was trained to discriminate rule identity (ABCD vs ABBA) using trials pooled across both pitch exemplars. The rule axis was fit using population activity in a late window spanning pips 3–4.

Because the number of units often exceeded the number of trials per class, we used a diagonal-covariance LDA (equivalent to Fisher LDA with a diagonal covariance estimate). Let 𝜇_%_and 𝜇_&_denote the class-conditional mean population vectors and 𝑣 the per-neuron variance (estimated on training trials). The discriminant weight vector was

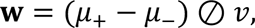

and test-trial scores were computed as 𝑠 = 𝐱𝐰 + 𝑏, where 𝑏 is the standard LDA intercept term computed from class means. Because LDA axes have an arbitrary sign, we aligned axis direction across cross-validation splits such that the mean projection of the positive class exceeded that of the negative class (high > low for pitch; ABBA > ABCD for rule). This ensured consistent interpretation of projection sign across sessions.

### Cross-validation and time-resolved decoding on fixed axes

All decoding performance was evaluated on held-out test data. Within each session, trials were split into train and test sets (50/50) independently for each condition, and the procedure was repeated across multiple random resamples (at least 20). For each resample we:

1. Fit the pitch axis using only training trials in the pitch training window.
2. Fit the rule axis using only training trials in the rule training window.
3. Applied these fixed axes to the held-out test trials at each time point to obtain scalar scores for each trial.

At each time point we quantified discrimination using the area under the ROC curve (AUC), computed from the distribution of test scores for the two classes (chance = 0.5). We computed four time-resolved AUC curves per session:

- **Pitch on PitchAxis:** AUC for low vs high pitch using projections onto the pitch axis.
- **Rule on RuleAxis:** AUC for ABCD vs ABBA using projections onto the rule axis.
- **Pitch leakage on RuleAxis:** AUC for low vs high pitch using projections onto the rule axis.
- **Rule leakage on PitchAxis:** AUC for ABCD vs ABBA using projections onto the pitch axis.

Leakage curves provide internal controls for axis specificity: if an axis is selective for one variable, decoding the other variable on that axis should remain at chance.

### Cross-condition generalisation performance (CCGP)

To quantify whether abstract rule information generalized across sensory instantiations, we computed cross-condition generalisation performance (CCGP) in the equiprobable 2×2 design (A_0_B_0_C_0_D_0_, A₁B₁C₁D₁, A_0_B_0_B_0_A_0_, A₁B₁B₁A₁). For each session, we formed trial-wise population response vectors by averaging neural activity across a short sliding integration window at each time point (relative to sequence onset), after optional baseline subtraction using a pre-sequence baseline. CCGP was evaluated using a strict cross-generalisation scheme: at each time point we trained a linear classifier to discriminate rule (ABCD vs ABBA) within one pitch band (train: A_0_B_0_C_0_D_0_, vs A_0_B_0_B_0_A_0_) and tested on the other pitch band (test: A₁B₁C₁D vs A₁B₁B₁A₁), and vice versa (train on band 1, test on band 0); performance was averaged across the two directions. Trials were randomly split into train/test sets within each condition (50/50, repeated across resamples). Neural features were standardized using training-set statistics only (per-neuron z-scoring computed on training trials and applied unchanged to test trials), and units with near-zero training variance were excluded. Decoder performance was quantified primarily by area under the ROC curve (AUC; chance = 0.5). Time-resolved CCGP curves were summarized across sessions as mean ± SEM. Statistical significance was assessed using permutation procedures that preserved the train/test structure (shuffling training labels, refitting the classifier, and evaluating on the held-out test set); for time-resolved inference, permutation-based cluster correction was used to control for multiple comparisons across time.

## Supplementary figures

**Supplementary Figure 1.**
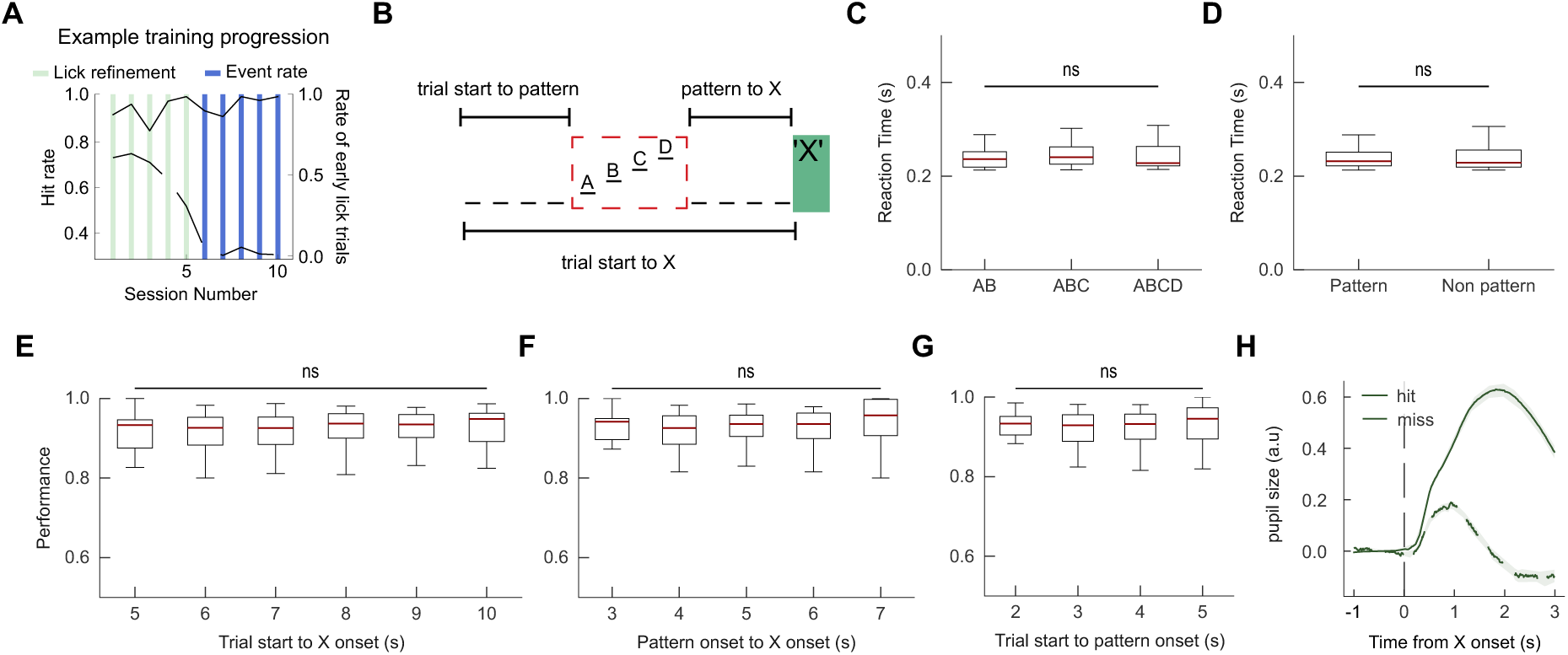
Behaviour across trial conditions. (A) Training progression for an example mouse over first 10 training sessions. Solid line: hit rate. Dashed line: proportion of trials with pre-X licking. Coloured bars mark training stages. Lick-refinement stage: mouse learned to lick only at the X stimulus (no sequences presented). Event-rate sessions: presentation rate of the pattern sequence was varied. (B) Schematic illustrating the temporal windows used for behavioural and pupil analyses. (C–H) Boxplots show performance metrics averaged across mice (n = 18) (red line, median; boxes, interquartile range; whiskers, 1.5 × IQR). (C) Reaction times for catch trials in which the X stimulus appeared during or immediately after the pattern sequence. (D) Reaction times on pattern versus non-pattern trials. (E) Hit rate as a function of trial start–to–X onset interval. (F) Hit rate as a function of pattern onset–to–X onset interval. (G) Hit rate as a function of trial start–to–pattern onset interval. (H) Pupil dilation response to X presentation on hit and miss trials (bold line, mean across mice; shaded area, SEM).

**Supplementary Figure 2.**
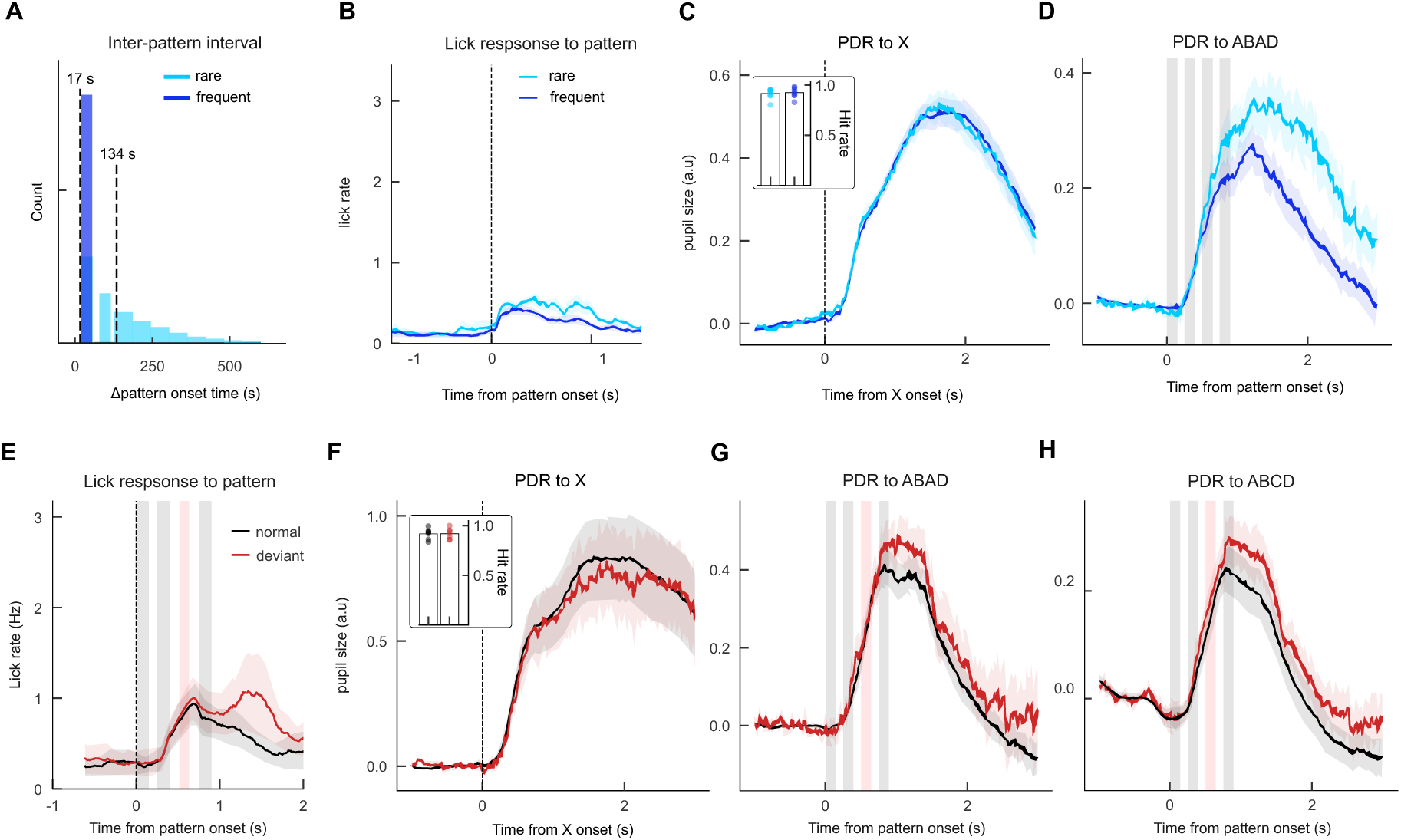
Pupil dilation response across conditions: (A) Distribution of inter-pattern intervals for rare and frequent trials across all sessions (n = 23304 frequent, 12305 rare trials from 17 mice). (B) Lick responses aligned to pattern onset for rare and frequent trials (bold line, mean across sessions (n = 413 sessions from 17 mice); shaded area, SEM). (C) Pupil dilation response (PDR) to the X stimulus (n = 223 sessions from 8 mice). Inset panel shows mean hit-rate across rare and frequent sessions averaged across mice (n = 8). No significant difference between performance on rare vs frequent trials (t(14): p = 0.74). (D) PDRs for rare and frequent trials in sessions where the pattern sequence was ABAD (n = 74 sessions from 9 mice). Dots show mean hit-rate for individual mouse. (E–H) Lick and pupil responses for normal and deviant pattern sequences. (E) Lick response averaged across sessions (142 sessions from 16 mice) (Vertical dashed line indicate sound event onsets (sequence in E and X in F). Vertical red bar denotes deviant tone presentation period. For all traces, bold lines represent the mean across sessions and shaded regions denote SEM. (F) PDR to X on normal vs deviant trials averaged across control sessions (n = from animals). Inset panel shows mean hit-rate across normal vs deviant sessions averaged across sessions (n = 28 from 7 mice). No significant difference between performance on normal vs deviant trials (t(12): p = 0.99). Dots show mean hit-rate for individual mouse. (G) PDR to pattern on normal and deviant trials averaged across sessions where normal pattern was ABAD and deviant pattern was ABCD (n = 21 sessions from 6 mice) (H) PDR to pattern on normal and deviant trials averaged across sessions (n = 106 sessions from 6 mice).

**Supplementary Figure 3.**
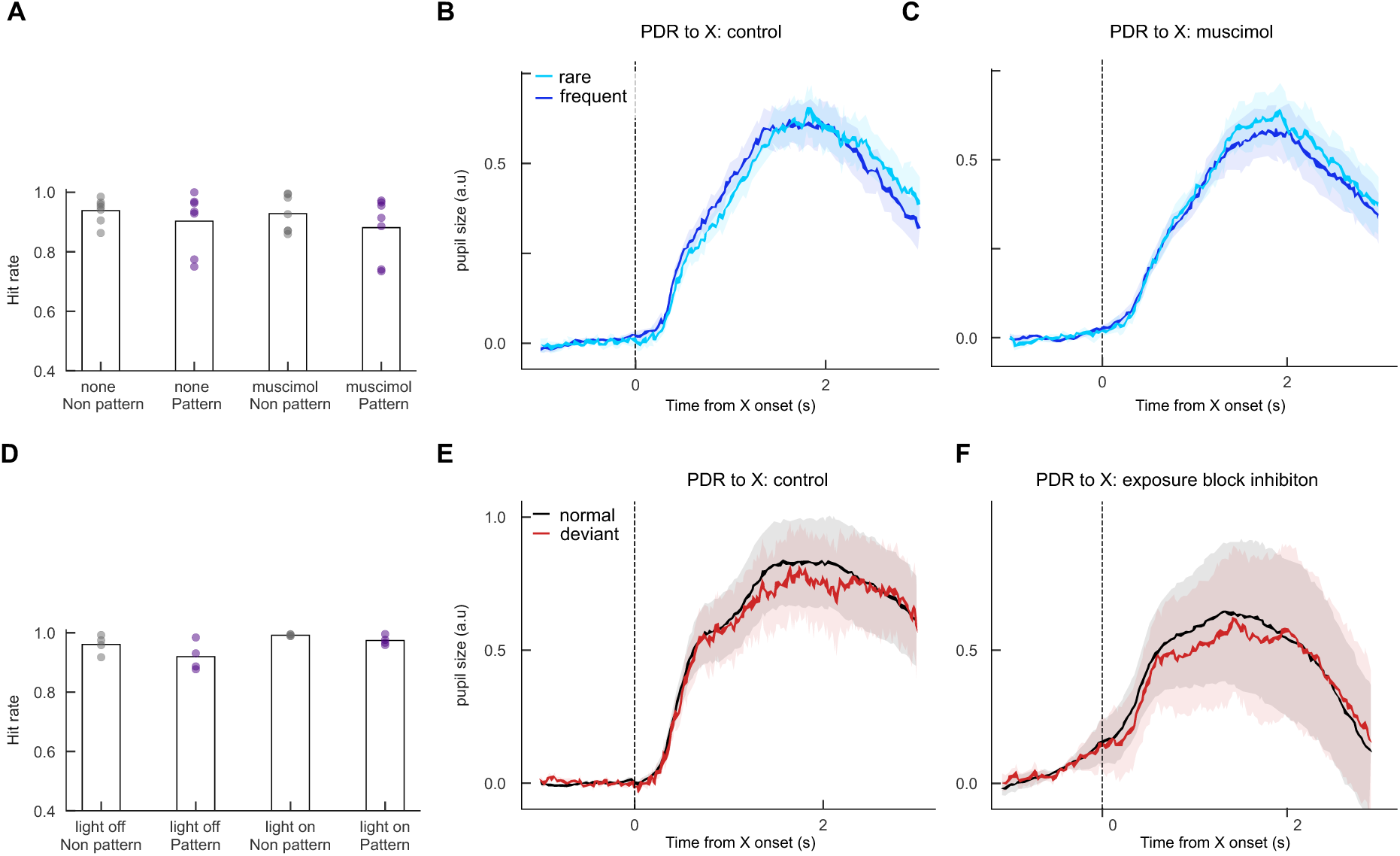
Behavioural performance and pupil responses during hippocampal inactivation. (A) Hit rate across non-pattern and pattern trials during control (none) and muscimol sessions. Bars show mean across mice (n = 6). Individual dots show mean across individual mice. (B–C) Pupil dilation responses (PDRs) aligned to X onset for non-pattern (B) and pattern (C) trials during control (dark blue) and muscimol (light blue) sessions. (D) Hit rate across non-pattern and pattern trials during optogenetic control (light off) and inhibition (light on) sessions. Bars show mean across mice (n = 4). Individual dots show mean across individual mice. (E–F) Corresponding PDRs for non-pattern (E) and pattern (F) trials during light-off (dark red) and light-on (light red) sessions. Bold lines indicate mean across mice; shaded regions denote SEM.

**Supplementary Figure 4.**
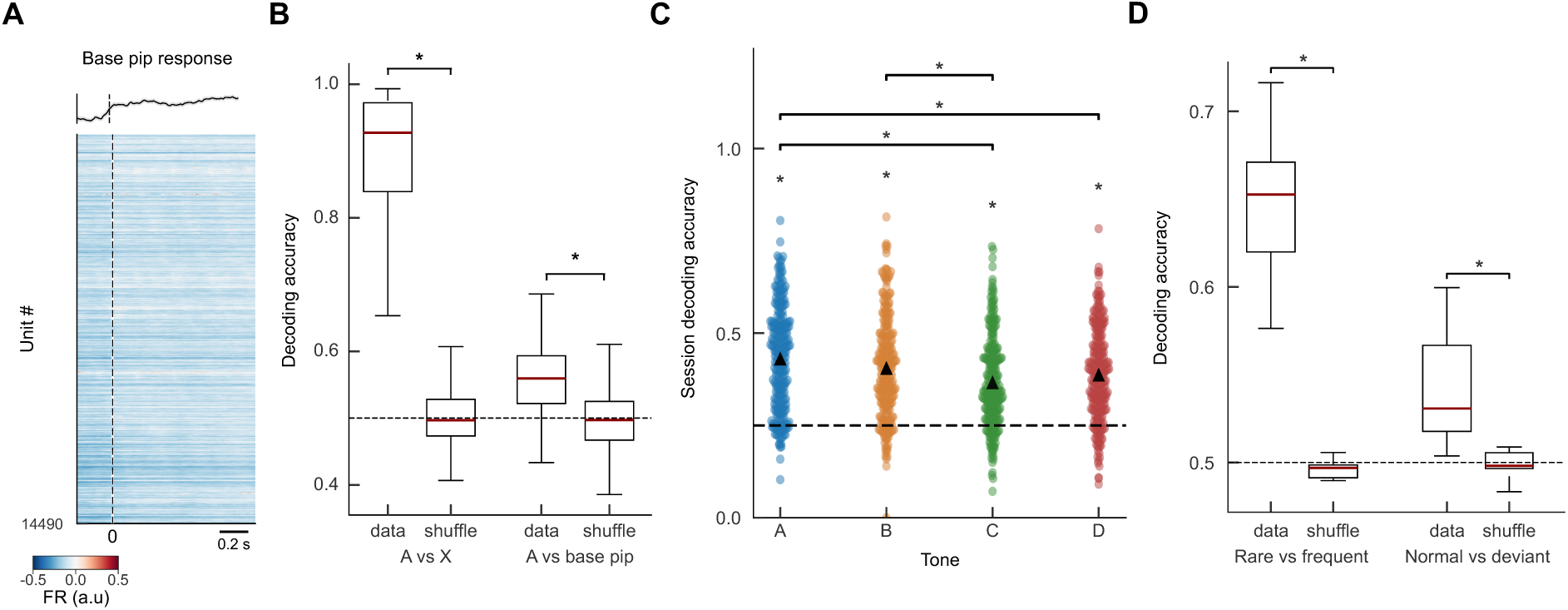
Neural decoding performance across sessions and conditions. (A) Population peristimulus time histogram (PSTH) aligned to tone onset for base pip, averaged across units and sessions. Top subpanel shows mean firing rate aligned to tone onset for base pip. averaged across units. (B) Decoding accuracy across sessions (n = 264 sessions from 6 mice) for A vs. X and A vs. base pip comparisons, showing performance for real data and label-shuffled controls. (C) Decoding accuracy for individual tone identities (A, B, C, D) across sessions. Circles show mean decoding accuracy for each session. Black triangles show mean across all sessions (n = 264 from 6 mice). Bonferroni post hoc and t-tests, as appropriate *P < 0.05. (D) Decoding accuracy averaged across mice for rare vs. frequent (n = 6) and normal vs. deviant (n = 4) comparisons. In boxplots, red lines indicate medians, boxes show interquartile ranges, and whiskers denote 1.5 × IQR.

**Supplementary Figure 5.**
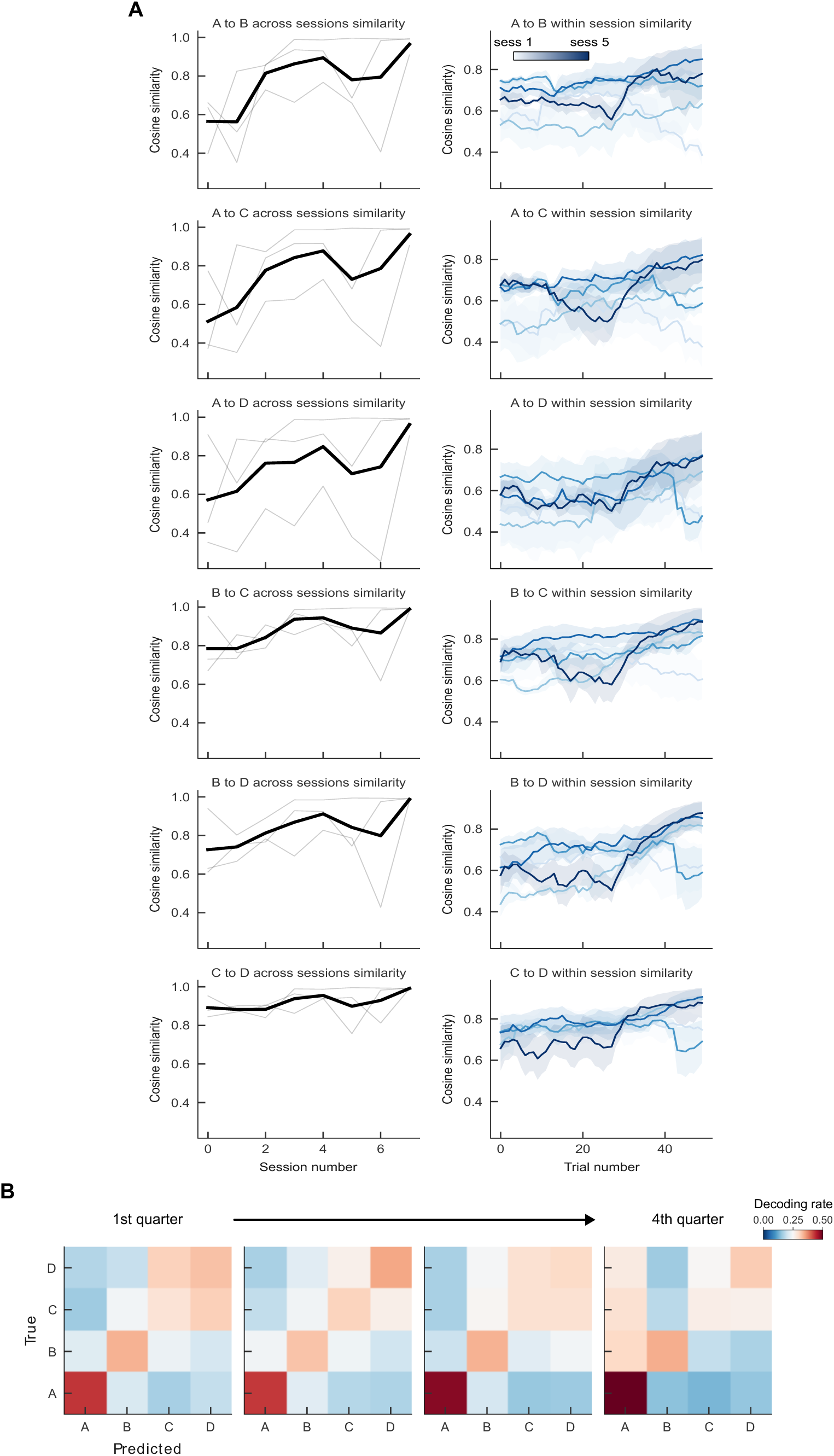
Evolution of tone representation similarity during early exposure to the pattern sequence. (A) Cosine similarity between tone-evoked population responses over the initial exposure sessions. Left: mean cosine similarity across the first five sessions (bold line, mean; light lines, individual mice; n = 3). Right: within- session similarity computed using a sliding 25-trial window. Shaded areas indicate SEM across mice (n = 3). (B) Stimulus decoding across the first five sessions. Confusion matrices show decoding accuracy for tones A, B, C, and D, separated into four temporal quarters within each session and averaged across the first 5 from 3 mice (n = 15).

**Supplementary Figure 6.**
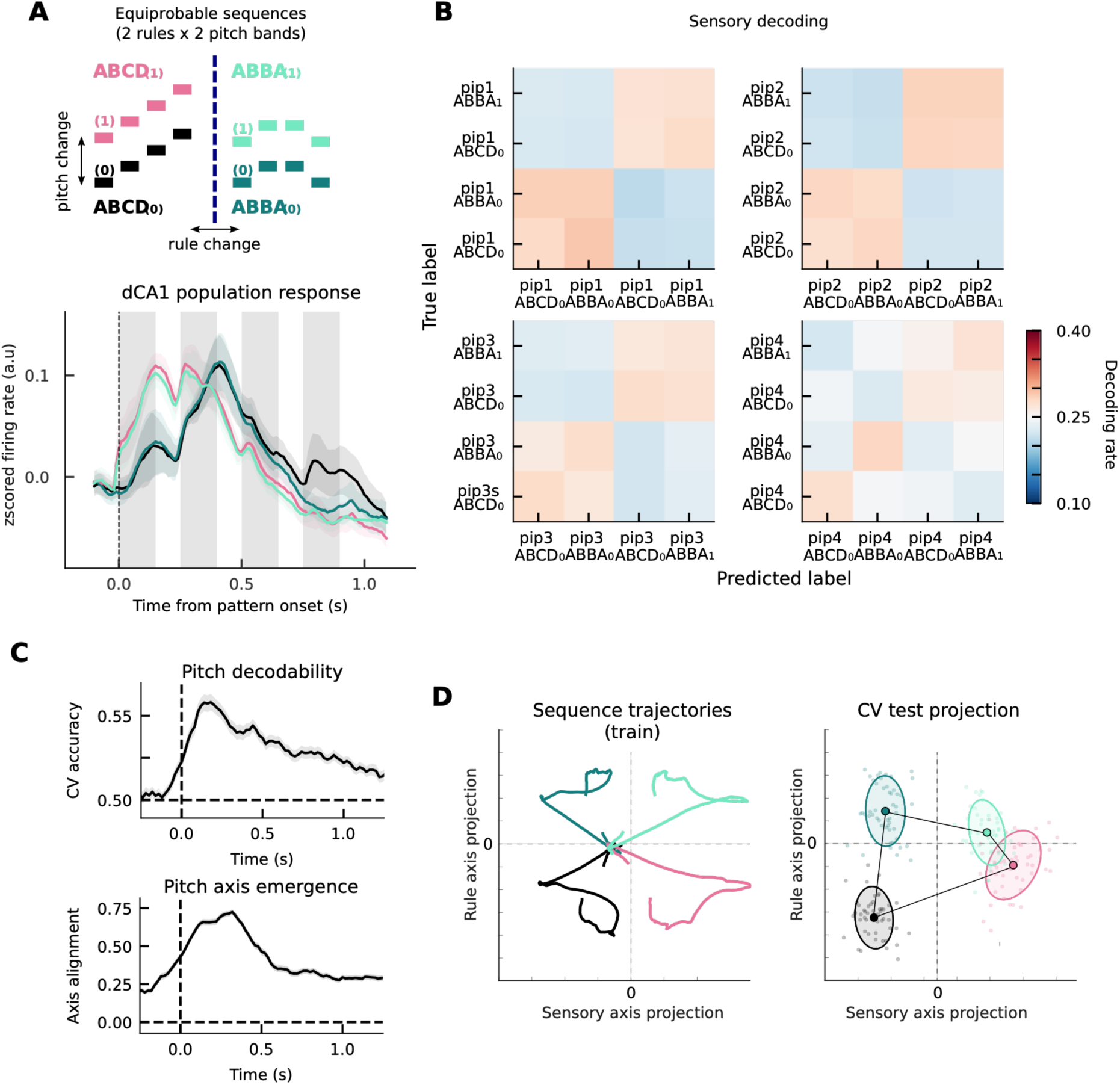
(A) Population response. Mean z-scored firing rates aligned to pattern onset for normal and deviant sequences. Shaded regions indicate SEM across mice; grey bars denote tone presentation windows. **(B) Pip decoding.** Confusion matrix for four-way classification of tones from 4 sequences, at each pip, averaged across sessions. **(C) Pitch axis emergence.** Top: Cross-validated LDA decoding accuracy for pitch identity computed in sliding time windows (chance = 0.5, dashed line). Bottom: Pitch axis emergence, quantified as the cosine alignment between LDA axes trained at each time window (sliding 200 ms window) and an early reference pitch axis (trained on pips 1–2). Dashed vertical line indicates pip1. **(D) The geometry of abstraction.** Left: State-space trajectories in the 2D plane defined by the pitch and rule axes), revealing a factorised “rectangle-like” geometry in which pitch and rule act as approximately separable dimensions of the population code. Right: Projection of pip 1-2 on Pitch axis, and pip 3-4 on Rule axis. Dots show individual sessions (n = 136, CV test trials).

**Supplementary Figure 7:**
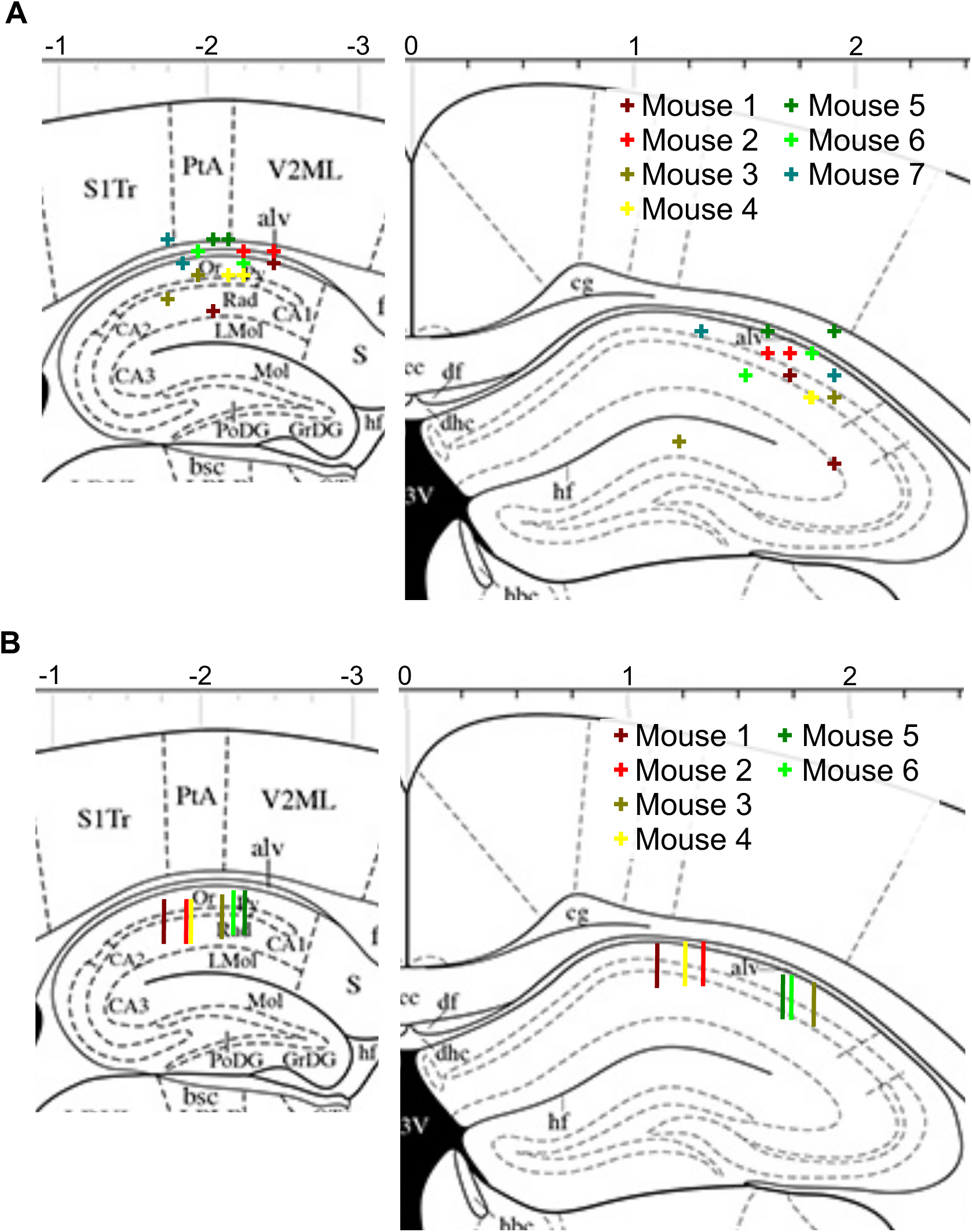
Anatomical locations for cannula infusion and silicon probe implantation. A) Diagram showing cannula locations estimated post-mortem. Positions corresponding to the centre of the infusion cannula plotted one sagittal (left) and coronal (right) hippocampal views. B) Diagram showing silicon probe locations estimated post-mortem. Positions corresponding to the centre of the 4-shank probe plotted one sagittal (left) and coronal (right) hippocampal views.

## Acknowledgment

The authors thank S. Hofer, T. Behrens, A. Macaskill and J. Harris for comments on the manuscript; the staff of the SWC Neurobiological Research Facility for animal support; the SWC Fabrication Laboratory for assistance with machining; and T. George for early help with the human version of the task. We also thank members of the Akrami lab and other SWC laboratories for insightful discussions and advice. This work was supported by core funding to the Sainsbury Wellcome Centre from Wellcome (219627/Z/19/Z) and the Gatsby Charitable Foundation (GAT3755), and by UK Research and Innovation (EP/Z000599/1) to A.A.

## Author information

Sainsbury Wellcome Centre for Neural Circuits and Behaviour, University College London, Howland St, London Adedamola Onih, Xinran Shen, Lida Pentousi, Vezha Boboeva, Athena Akrami

## Contribution

A.O. and A.A. conceived the project. A.O. carried out all experiments and analysed the data, with assistance from L.P., X.Sh., A.A. and V.B.. A.O. and A.A. wrote the manuscript with comments from V.B.

